# An organ-resolved rat FFPE phosphoproteome map enables directional kinase activity inference

**DOI:** 10.64898/2026.07.28.741173

**Authors:** Erin M Humphries, Marius Schliemann, Naomi O’Sullivan, Peter G Hains, Phillip J Robinson, Bernhard Kuster

## Abstract

Formalin-fixed paraffin-embedded (FFPE) tissue is the dominant clinical pathology resource yet whether it faithfully preserves organ signalling biology and supports directional regulatory analysis remains unquantified. We generated a phosphoproteome map from eight healthy rat organs, separating preservation effects from biological variation. Using mass spectrometry, we quantified 54,710 phosphosites on 5,994 proteins across receptors, kinase cascades and nuclear regulators. Organ-specific phosphosite signatures matched known physiological and proliferative states. Paired antagonistic phosphosites converted into “activating-minus-inhibitory” indices that quantified net tissue-specific pathway activity, while a “kinase-by-organ activity” matrix resolved functional hierarchies.

Joint analysis with an external fresh-frozen phosphoproteome dataset yielded 58,631 phosphosites total, recovering 86% of the 28,888 sites detected in the frozen dataset. Organ identity explained over 92% of the total variance after batch correction, versus under 0.5% for preservation method. Per-organ phosphosite intensities agreed closely between preservation modes except in brain. This establishes that archived pathology tissue supports biologically faithful phosphoproteome analysis at organ, pathway, and site resolution, providing a framework for retrospective signalling studies in clinical archives.

**Graphical Abstract:** 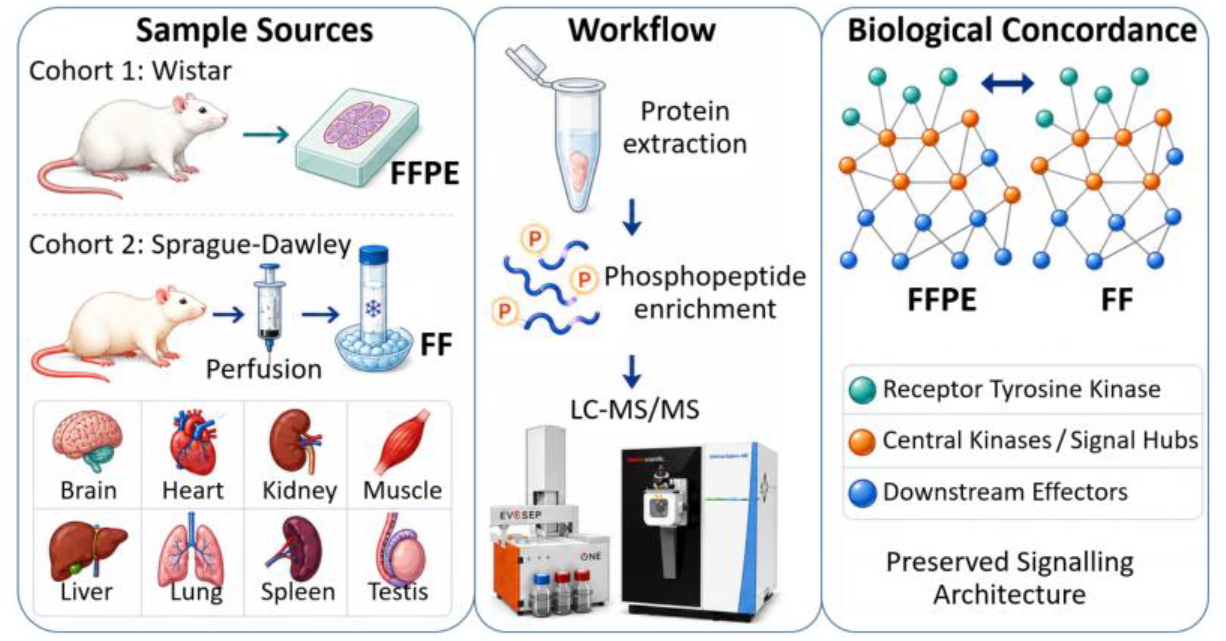

## Introduction

Protein phosphorylation is a central regulatory layer controlling cellular proliferation, metabolism and survival. While genomic alterations indicate signalling potential, quantitative phosphosite profiles report functional pathway activity and targetable dependencies that remain undetectable by protein-abundance measurements alone.^1, 2^ Translating this systems-level information into clinical practice requires phosphoproteomic measurements that are deep and quantitatively stable in genuine clinical specimens. Formalin-fixed paraffin-embedded (FFPE) tissues represent the largest and most clinically relevant resource for translational oncology and biomarker discovery, as they are generated routinely in pathology and stored in massive, clinically annotated archives.

Early mass spectrometry-based analyses of FFPE tissue were hindered by formalin-induced protein crosslinking. However, advances in sample preparation have established the feasibility of deep proteomic and phosphoproteomic profiling from archival material.^3–32^ Recent liquid chromatography-tandem mass spectrometry (LC-MS/MS) studies using data-independent acquisition (DIA) have further reduced stochastic MS2 sampling, enabling consistent quantification of thousands of phosphosites.^33^ Despite these technical advances, the FFPE phosphoproteome is necessarily influenced by pre-analytical variables. Phosphorylation networks are highly dynamic; specific sites may be lost during cold ischemia, stress pathways can activate during the delay to fixation, and matrix effects may alter extraction efficiency compared to fresh-frozen (FF) tissue.^34–43^

Phosphoproteomics of FF tissue is moving beyond biological characterisation into clinical decision-making. A recent pan-cancer precision oncology study profiled proteomes and phosphoproteomes from nearly 2,000 tumour samples across real-world molecular tumour board programs, deriving kinase activity scores and immune signalling readouts that informed treatment recommendations beyond what genomic sequencing alone could capture, including stratifying drug response in sarcoma and chordoma.^44^ A companion platform paper describes the infrastructure built to convert this phosphoproteomic data into patient-specific reports usable at the point of clinical care.^45^ Together, these studies show that phosphoproteomic signalling readouts, including kinase activity scores derived from targeted substrate panels, can be measured at scale and carry direct clinical utility, but they currently rely on FF material, which is far less available than the archival FFPE specimens that dominate pathology biobanks. Yet, a critical methodological and biological gap remains: the fidelity of FFPE phosphoproteomes relative to FF is poorly defined. It is currently unclear which components of organ-specific signalling are preserved in FFPE and which are distorted by pre-analytical effects. Furthermore, while technical benchmarking studies have compared extraction efficiencies, the field lacks a multi-organ assessment demonstrating whether FFPE phosphosite intensities retain enough biological structure to resolve antagonistic regulation or directional signalling. It similarly remains unclear whether FFPE preserves quantitative kinase substrate activity at a level comparable to FF tissue. If FFPE phosphoproteomes could be shown to retain the same signalling structure that underlies these FF-based clinical tools, the enormous existing archive of banked FFPE specimens could be brought into this framework. Whether that structural fidelity holds, however, has not been systematically tested.

To address this gap, we generated a high-depth DIA-MS FFPE phosphoproteome map across eight healthy rat organs. We applied this map to determine whether FFPE-derived phosphoproteomes recapitulate expected organ physiology from receptor cascades to chromatin regulators. We subsequently quantified whether FFPE phosphosite intensities can resolve paired activating and inhibitory regulatory sites to yield directional kinase activity scores. Finally, we integrated this FFPE map with an external multi-organ FF phosphoproteome dataset to quantify preservation-related matrix effects. As a healthy-tissue reference map, this is a proof-of-concept study that demonstrates FFPE phosphoproteomes can be used for retrospective interpretation of signalling states in archival pathology cohorts.

## Methods

### FFPE Cohort and Experimental Design

The FFPE cohort comprised 74 LC-MS/MS raw files generated from eight healthy Wistar rat organs (brain, heart, kidney, liver, lung, leg muscle, spleen, testis). For each organ, one bulk protein digest was prepared per rat and phosphopeptides were enriched from replicate aliquots. Technical replicates were defined as independent phosphopeptide enrichments from the same digest: five enrichments (50 μg peptide input each) were performed per organ (N = 4 for spleen), producing 39 FFPE raw files. All five technical enrichments were performed from a single rat digest per organ to ensure that between-replicate variance reflects enrichment reproducibility rather than biological heterogeneity. Biological replicates were defined as distinct rat–organ combinations. For kidney, liver, and testis, up to four enrichments per organ were acquired from each of three rats, yielding an additional 35 FFPE raw files and providing multi-animal coverage for these organs. Sample annotations and MS files are detailed in **Supplementary Table 1.**

### Sample Preparation

FFPE organs from adult male Wistar rats (6–8 weeks, 250 g) were processed into FFPE blocks as described.^6, 32^ Eight sequential 60 μm sections per block were cut using a motorised rotary microtome, pooled, and stored at −20°C for <1 week. Tissues were deparaffinised with heptane/methanol, lysed in 4% SDS/500 mM Tris-HCl (pH 9.0), homogenised (tissue lyser, 2 min, 200 Hz), sonicated (Covaris R230, 5 min), and heated (90°C, 10 min).^11, 14^ SDS was removed by SP3 clean-up with magnetic beads (Sera-Mag Speed Beads A and B, in a 1:1 ratio) and 70% ethanol precipitation.^46^ Bead-bound proteins were reduced (20 mM DTT, 45 min), alkylated (55 mM chloroacetamide, 30 min), and digested overnight with trypsin (1:100 ratio, assuming 600 μg total protein per sample) in 40 mM Tris-HCl (pH 7.8) supplemented with 2 mM CaCl_2_ at 37°C. Peptides were recovered, acidified, and desalted using C18 Sep-Pak cartridges (50 mg). Peptide concentration was determined by UV absorbance (280 nm).

Phosphopeptides were enriched using mixed Ti-IMAC HP and Zr-IMAC HP beads (2 μL each, ReSyn BioSciences) in 96-well format.^47^ Beads were incubated with 50 μg peptide (30 min, RT), washed sequentially with 80% acetonitrile/1% trifluoroacetic acid followed by 10% acetonitrile/0.2% trifluoroacetic acid, eluted twice with 1% ammonium hydroxide, and loaded onto Evotips (EvoSep Pure, EV2011).^14^ For the global proteome analysis, approximately 1 µg of peptide digest was loaded onto Evotips.

### LC-MS/MS

The FFPE cohort was analysed on an Evosep One LC system (EvoSep Biosystems) coupled to an Orbitrap Exploris 480 mass spectrometer (Thermo Fisher Scientific) with an Endurance column (15 cm × 150 μm, 1.9 μm C18, room temperature) with the 15 SPD method (220 nL/min, 88 minute gradient).^6^ Data acquisition comprised a full MS1 scan (360–1000 m/z, 60,000 resolution, AGC 300%, 100 ms injection time) followed by 42 DIA windows (15 m/z width with 1 m/z overlap, HCD 28%, 30,000 resolution, AGC 1000%, 50 ms injection time, 2.2 s total cycle time).

### FF Cohort and Experimental Design

The FF cohort comprised three Sprague–Dawley biological replicates per organ (24 raw files total) from a published dataset (PMID: 32051234).^48^ Rats were perfused, organs snap-frozen, and processed by automated PAC digestion, Ti-IMAC HP enrichment, and Evotip loading. FF samples were analysed on an Evosep One–Orbitrap Exploris 480 using an in-house column (15 cm × 150 μm, 1.9 μm C18, 60 °C) with the 60 SPD method (1 μL/min, 21-min gradient). MS1 scans were acquired at 120,000 resolution (45 ms injection time) followed by 49 DIA windows (13.7 *m/z* width with 1 *m/z* overlap, 15,000 resolution, 22 ms injection time; 1.1 s total cycle time).

### Data Searching

DIA files were analysed in Spectronaut (version 19.7) using the directDIA+ (Deep) workflow.^49^ The UniProt *Rattus norvegicus* database (Swiss-Prot and TrEMBL, 92,920 sequences; downloaded 22 January 2025) was searched with Trypsin/P and LysC/P specificity, ≤ 2 missed cleavages, peptide lengths between 7-52 amino acids, and carbamidomethylation of cysteine as a fixed modification.^50^ Variable modifications included methionine oxidation, protein N-terminal acetylation and serine/threonine/tyrosine phosphorylation (maximum five per peptide). False discovery rate was controlled at 1% at the PSM, peptide and protein levels using target-decoy strategy with pseudo-reverse (“KR”) decoys. Quantification used the Biognosys Phospho PTM Workflow, with cross-run disabled and a phosphorylation sites localisation probability of at least 0.75. This threshold was selected to maximise site coverage for map-level analyses while retaining sites with a confidence level consistent with published FF phosphoproteomics benchmarks.

### Data Processing and Phosphosite Consolidation

All processing was performed in R (v4.5.1) with fixed seed (123) and L’Ecuyer-CMRG generator for reproducibility.^51^ The Spectronaut PTM site report was filtered to retain only phosphorylated precursors. Each phosphosite received a unique identified (format Protein_Site_Gene, e.g., P13596_S774_Ncam1). When multiple protein shared the same phosphorylation site (identical PTM flanking sequence), SwissProt-entries (prefix P/Q/O) were prioritised over TrEMBL, and redundant entries removed.

### FFPE Phosphoproteome Map and Regulatory Phosphosite Heatmap

A curated panel of phosphosites implicated in growth factor, stress, immune, developmental, cell-cycle, and DNA damage pathways was assembled for organ-level visualisation. Per-site organ means were calculated by averaging normalised log₂ values across all samples from the same organ, then z-scored per site by subtracting the across-organ mean and dividing by the across-organ standard deviation; non-variable sites were assigned a z-score of zero. Positive values indicate above-average phosphorylation relative to a site’s across-organ mean; negative values indicate below-average phosphorylation.

### Antagonistic regulation

To quantify directional net signalling activity at key regulatory nodes, we selected four proteins for which antagonistic phosphosite pairs are well established in the literature as markers of activation versus inhibition, and for which both sites were reliably quantified across all eight organs in our dataset: Gsk3a (activating Y279, inhibitory S21), Ctnnb1 (activating S552, inhibitory S45), Cdk1 (activating T161, inhibitory T14), and Bad (pro-survival S167, pro-apoptotic Y114). These four were prioritised as representative case studies spanning distinct regulatory contexts, metabolic kinase signalling, transcriptional co-activation, cell-cycle control, and apoptotic balance, rather than as an exhaustive survey, because consistent, high-confidence quantification of both sites in a pair across all organs was required to compute a reliable index. For each protein, mean normalised log₂ intensity was computed per site per organ across all samples without further scaling. Activity indices were calculated per organ as the difference between the activating and inhibitory site (activating minus inhibitory). For Bad, the index was defined as S167 minus Y114, where positive values denote a net pro-survival phosphorylation bias and negative values denote a net pro-apoptotic bias. For kinase regulators (Gsk3a, Cdk1) and the transcriptional co-activator Ctnnb1, positive index values denote net activating phosphorylation, and negative values denote net inhibitory phosphorylation. These indices are intended as rank-order comparisons of relative net activation state across organs and do not report absolute enzymatic output.

### PI3K–AKT routing panels

Functional site panels representing upstream Pi3k-Akt activation and major downstream branches were defined: Mtor/metabolism, translation, acetyl-CoA carboxylase (ACC), survival/adaptors, junctions, and adhesion. For each phosphosite, mean log₂ intensities of median-normalised values were first computed per organ. Each phosphosite was then z-scored once, across all eight organs, such that every single site has a single mean-zero, unit-variance distribution spanning the full organ panel. This ensures that all z-scores entering the averaging step are computed from the same underlying distribution and are therefore directly comparable across organs. Route indices were derived by averaging these per-organ z-scores across all sites within each functional group, yielding an upstream Akt panel (Pdpk1 S244, Akt1 S124, Akt2 S474, Aktip S30) and downstream panels: mTOR/metabolism (Mtor S1230, Rptor S722, Mapkap1 T462, Ahcy Y7, Bckdha Y346, Akt1s1 T247), translation (Eif4b S406/S422, Eif4e S209, Eif4ebp1 T76, Eif4g1 S1130, Eif4g2 T446), ACC (Acaca S79, Acacb S219), survival/adaptors (Bad Y114/S167, Gab1 Y259, Shc1 S29, Ptpn11 Y605), junctions (Tjp1 Y895, Tjp2 Y1155, Pxn S83), and adhesion (Ctnnb1 S552, Ctnnd1 S268, Apc Y2837, Axin1 Y82, Ptk2 Y576, Gsk3a Y279/S21, Gsk3b T7/S9). ACC–Akt coupling was assessed by regressing the ACC route index on the upstream Akt index; residuals were retained as an ACC uncoupling score (observed minus expected ACC activation) per organ.

### Kinase Substrate Activity Scores

Phosphosite intensities were normalised against a matched global proteome dataset. This step separates two sources of signal that are otherwise confounded: an increased in phosphosite intensity driven by increased kinase activity, versus an increase driven simply by higher abundance of the parent protein in that organ. Mean log2 protein group intensity per organ was used as the normalisation baseline. Normalisation followed one of two rules, depending on whether the parent protein was detected in the proteome dataset. Where the protein was detected, proteome log2 intensity was subtracted from phosphosite log2 intensity. Where the protein was not detected, protein abundance was assumed constant across organs, and the raw phosphosite intensity was retained unadjusted.

These two rules produce log2 intensity values on different numerical scales. Phosphosites with a measured parent protein yield subtracted values that cluster around zero, whereas phosphosites without a measured parent protein retain the full magnitude of the raw log2 scale. To correct this scale mismatch, each phosphosite was z-scored individually across the eight organs, irrespective of whether its parent protein had been detected in the proteome dataset. Because each phosphosite is scored only against its own across-organ values, the phosphosite’s original absolute scale is removed entirely, leaving only its relative pattern: which organs show higher or lower phosphorylation for that specific phosphosites. The conditional subtraction rule is a conservative choice. It may slightly overestimate phosphorylation stoichiometry for proteins whose organ-specific abundance differences fall below the detection limit of the proteome dataset.

Kinase activity was inferred using curated kinase–substrate relationships established via potency-coherence analysis, in which dose-response curves for 133 kinase inhibitors with known targets and affinities were measured across the phosphoproteome.^52^ Phosphosites were assigned to a kinase when their inhibitor sensitivity profile was quantitatively coherent with that kinases’ known inhibitor potency profile (icorr > 0.8, adjusted *p* < 0.05). For each of 28 kinase families, we took all of that kinases’s known substrate phosphosites (already z-scored), median centred them per sample, and summed them into one activity score per kinase family per organ. Each kinase family’s activity score was then z-scored a second time, this time across the eight organs for that kinase family specifically, so that each of the 28 kinase families has its own across-organ z-score distribution. Organ-level means and standard errors of these per-kinase, across-organ z-scores were then calculated.

Muscle and heart tissue contain very high amounts of a small number of structural proteins, such as actin and myosin. Because these proteins make up a large share of the total proteome in these two organs, they distort the proteome baseline used in the normalisation step above, which can make proteome-normalised ratios appear artificially high in muscle and heart specifically, independent of any real kinase activity. Because the z-scoring above was performed per kinase family, across organs, and not per organ, organs are not centred at zero. Thus, a shared, organ-wide technical offset shared across kinase families can still be present within each organ. To remove this artefact, the median z-score across all 28 kinase families was calculated within each organ and subtracted from every individual kinase family’s zscore in that organ. This removes a shared, organ-wide technical offset that is unrelated to any single kinase’s true activity. Corrected values were then re-z-scored across organs a final time, producing the final between-organ kinase activity metric used in all downstream analyses.

### FFPE and FF Map integration

Phosphosite intensities from FFPE and FF samples were log₂-transformed and median-normalised. Missing values were addressed in four sequential steps: (i) organ-specific biological missingness was imputed using ptImpute (PhosR; percent1 = 0.5, percent2 = 0.1) applied pairwise across organs; (ii) phosphosites with less than 10% detection were removed; (iii) within-organ technical dropouts were modelled using scImpute (PhosR; threshold = 0.375); and (iv) remaining missing values were imputed by k-nearest neighbours (k = 10, rowmax = 0.99). The resulting matrix of 31,169 phosphosites comprised 70.1% measured and 29.9% imputed values. Systematic technical differences between preservation modes were removed using ComBat (sva v3.58.0; parametric priors, no covariates) applied to the complete post-imputation matrix.

### Phosphosite detection summaries and variance components

Detection summaries were generated from the pre-imputation, log₂ median-normalised matrix (58,631 sites). A phosphosite was considered detected if observed in at least one sample. Global FFPE–FF detection intersections were computed using VennDiagram (v1.7.3), and detected phosphosites were counted per organ and preservation mode. Within-organ, within-preservation-type reproducibility was summarised as the median Pearson r across all pairwise same-organ, same-preservation sample combinations on the batch-corrected matrix, restricted to natively detected values.

### Sample–Sample Correlation and Hierarchical Clustering

Pairwise Pearson correlation was performed on all FF–FFPE sample pairs per organ. To avoid artefactual inflation of correlation estimates, imputed values were masked prior to all calculations; correlations were therefore computed exclusively on natively detected, overlapping measurements using pairwise-complete observations (minimum 10 finite overlapping values per pair). This was performed independently on pre- and post-batch-correction matrices. Correlation distributions were visualised per organ as paired boxplots.

### FF–FFPE concordance

For organ-wise intensity comparison without imputation artefacts, the batch-corrected matrix with original missingness restored (imputed values masked) was used, restricted to phosphosites observed at least once per organ. Mean intensity per phosphosite was calculated separately across FF and FFPE samples per organ. A linear regression of FFPE on FF mean intensities was fitted per organ and visualised as faceted scatterplots annotated with the regression line, a 1:1 reference line, and the organ-specific equation and *R*².

## Data availability

The raw mass spectrometry data has been deposited in the ProteomeXchange Consortium via the PRIDE partner repository with the dataset identifier PXD057285.

## Results

In the sections below, we first characterise the FFPE phosphoproteome map to establish organ-specific phosphosite patterns. These patterns are subsequently validated through integration with an independent FF dataset in the final Results section.

### FFPE phosphoproteomes recapitulate organ-level signalling physiology

We first asked whether FFPE phosphoproteomes match expected organ physiology at phosphosite resolution. We generated an FFPE-only phosphoproteome map across eight healthy organs, enriching phosphopeptides from brain, heart, kidney, liver, lung, skeletal muscle, spleen, and testis and analysing them by LC-MS/MS.

Database searching against a merged Swiss-Prot and TrEMBL reference, included because the rat proteome remains incompletely annotated in Swiss-Prot alone, identified 90,255 phosphosites on 6,079 protein groups (**Supplementary Table S2**). After merging Swiss-Prot and TrEMBL entries, these collapsed to 54,710 sites on 5,994 protein groups corresponding to 5720 genes (**Supplementary Table S3**). This collapse was driven primarily by removal of redundant TrEMBL entries sharing identical PTM flanking regions with Swiss-Prot entries. The global phosphosite distribution was 74% phosphoserine, 21% phosphothreonine, and 5% phosphotyrosine.

FFPE phosphoproteomes resolved a multi-layered signalling architecture spanning receptor-proximal events, intracellular cascades, and nuclear control (**Figure 1**). This hierarchical coverage extended from ligand receptors to receptor tyrosine kinases, adaptor scaffolds, and cytoplasmic cascades (Raf–Mek–Erk, Pi3k–Akt–Mtor–S6k, Ampk, β-catenin, Hippo, Nf-κb/Tlr) to nuclear transcription factors. We identified 2,807 phosphotyrosine sites, including canonical activation-loop epitopes (e.g., Erk1/Mapk1 T183/Y185, Erk2/Mapk3 T203/Y205, Jnk1/Mapk8 Y185, and p38α/Mapk14 T170/Y172). Regulatory tyrosines were detected across additional signalling tiers (e.g., Gsk3a Y279, Fak/Ptk2 Y576, Shp2/Ptpn11 Y580).

**Figure 1.**
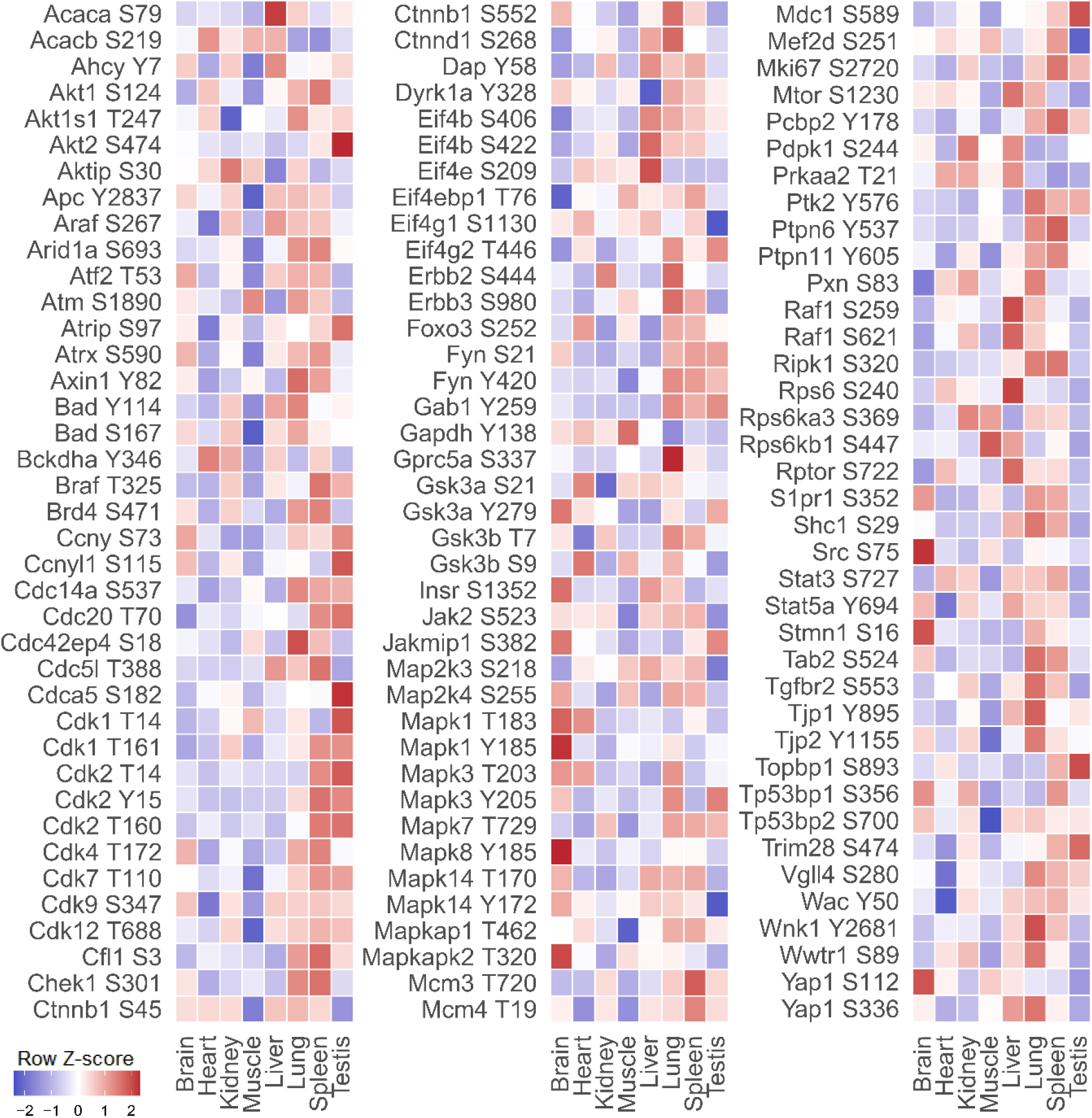
Body compartment distribution of key regulatory signalling networks across the rat FFPE phosphoproteome. The heatmap displays FFPE phosphosite intensities across eight healthy rat organs, highlighting core regulatory sites governing growth factor signalling, stress responses, immunity, development, and cell-cycle/DNA damage pathways. Phosphosite intensities are expressed as row-wise z-scores derived from median-normalised log2 intensities, representing the relative phosphorylation level of each site across organs without correcting for underlying protein abundance. Warmer and cooler colours indicate above- and below-average phosphorylation levels for a given site, respectively. Colour scale is clipped at ±2 z-score units to maximise dynamic range visibility.

### FFPE phosphoproteomes resolve organ-specific and cell-cycle signalling states

We next evaluated whether FFPE phosphoproteomes resolve tissue physiology and proliferative status. We assembled a curated panel of signalling sites spanning major pathways and quantified their phosphorylation across the eight organs (**Supplementary Table S4**). For this analysis, phosphosite intensities were expressed as non-proteome-normalised row *z*-scores, reflecting the relative phosphorylation of each site across organs without conflating differences in underlying protein abundance (**Figure 1**). These z-scores quantified relative phosphorylation enrichment per site across the organ panel.

FFPE phosphoproteomes showed reproducible, organ-specific phosphorylation patterns consistent with known tissue physiology. For metabolic and growth signalling, the map resolved high Mapk and Gsk3 activity in the brain (e.g., Erk1 T183/Y185, Erk2 T203/Y205, Gsk3a Y279). Heart tissue showed phosphorylation consistent with regulated fatty-acid metabolism and Gsk3-linked growth control (e.g., Acacb S219, Bckdha Y346, Gsk3a S21), whereas the kidney displayed Akt/Ampk and adhesion-associated signalling characteristic of transport epithelia (e.g., Aktip S30, Pdpk1 S244, Pxn S83). In the lung, epithelial survival and junctional regulators were preferentially phosphorylated (e.g., Bad Y114/S167, Ctnnb1 S552, Tjp1 Y895). This indicates that FFPE-derived phosphoproteomes capture the structured, tissue-resolved deployment of metabolic and growth pathways.

For proliferation and cell-cycle control, FFPE phosphoproteomes distinguished proliferative from largely post-mitotic tissues by jointly reporting cyclin-dependent kinase (Cdk) activation and replication machinery. The spleen and testis showed elevated phosphorylation of Cdk activation sites and proliferation markers (e.g., Cdk1 T14/T161, Cdk2 T14/T15, Mki67 S2720), consistent with active cell division in lymphoid and germ-cell compartments. The testis additionally showed phosphorylation consistent with Akt-linked survival, meiotic processes, and DNA-damage checkpoint control (e.g., Akt2 S474, Mdc1 S589, Trim28 S474). By contrast, brain, heart, and skeletal muscle displayed lower Cdk1/2 phosphorylation and reduced activity of transcriptional Cdks (e.g., Cdk7 T110, Cdk9 S347, Cdk12 T688). Skeletal muscle instead showed phosphorylation patterns consistent with metabolic regulation in post-mitotic tissue (e.g., Gapdh Y138, Rps6kb1 S447). Additional regulators further resolved tissue-specific organization of cyclin, cytoskeletal, and transcriptional modules (e.g., Ccnyl1 S115, Cdc42ep4 S18, Mef2d S251). FFPE phosphoproteomes thus capture both Cdk activation state and the partition between proliferative and post-mitotic signalling across tissues.

For stress, DNA-damage, and barrier functions, FFPE phosphoproteomes captured coordinated signalling consistent with tissue-specific demands. In the liver, we observed elevated phosphorylation of Raf–Mek–Erk and Akt–mTOR pathway components (e.g., Raf1 S259/S621, Pdpk1 S244, Rptor S722, Mtor S1230, Rps6 S240), consistent with metabolic activity and stress adaptation. In the lung, increased phosphorylation of epithelial survival, junctional, and Wnt regulators (e.g., Akt1s1 T247, Axin1 Y82, Bad Y114/S167, Ctnnb1 S552, Wwtr1 S89) was observed, consistent with barrier tissue signalling. DNA-damage response markers showed tissue-resolved phosphorylation patterns (e.g., Atm S1890, Atrip S97, Atrx S590), with downstream checkpoint effectors following the same organization (e.g., Chek1 S301, Mdc1 S589, Topbp1 S893). Replication-licensing and mitotic-exit markers were enriched in spleen and testis (e.g., Cdc20 T70, Cdca5 S182, Mcm3 T720), consistent with proliferative compartments. Across these examples, FFPE phosphoproteomes recover integrated, tissue-appropriate networks of DNA-damage response, survival signalling, barrier maintenance, and replication control.

### Antagonistic phosphosite pairs reveal directional signalling states in FFPE tissue

We then asked whether FFPE phosphoproteomes resolve antagonistic regulation on individual proteins by quantifying paired activating and inhibitory phosphosites across organs. We defined activating-minus-inhibitory indices using organ-level median log2 phosphosite intensities, where positive values indicate net activation and negative values indicate net inhibition. These indices provide a summary of relative net activation and should be interpreted as rank-order comparisons across organs rather than absolute measures of enzymatic output. Because the phosphosites used in these indices are a subset of those used in downstream kinase–substrate annotations, such indices directly inform inferred kinase substrate activity when aggregated across proteins.

To quantify net activity at metabolic and transcriptional hubs, we examined Gsk3a and Ctnnb1. Gsk3a Y279 activation was highest in brain and testis, whereas the inhibitory S21 site was most elevated in heart (**Figure 1**). The Y279–S21 index identified the highest net Gsk3a activity in brain, kidney, and testis, and lowest values in spleen, skeletal muscle, and liver, resolving tissue-specific differences in net kinase activation state (**Figure 2A**). For Ctnnb1, activating S552 and inhibitory S45 sites were both elevated in the lung, yet the S552–S45 index still ranked the lung (and to a lesser extent skeletal muscle) highest in net activity. This demonstrates that directional indices recover net signalling output even when activating and inhibitory inputs co-occur and simple phosphosite abundance would be ambiguous. For both proteins, activating-minus-inhibitory indices were elevated in tissues where these pathways are expected to be engaged, indicating that FFPE phosphoproteomes capture quantitative, directional pathway activity from antagonistic site pairs.

**Figure 2.**
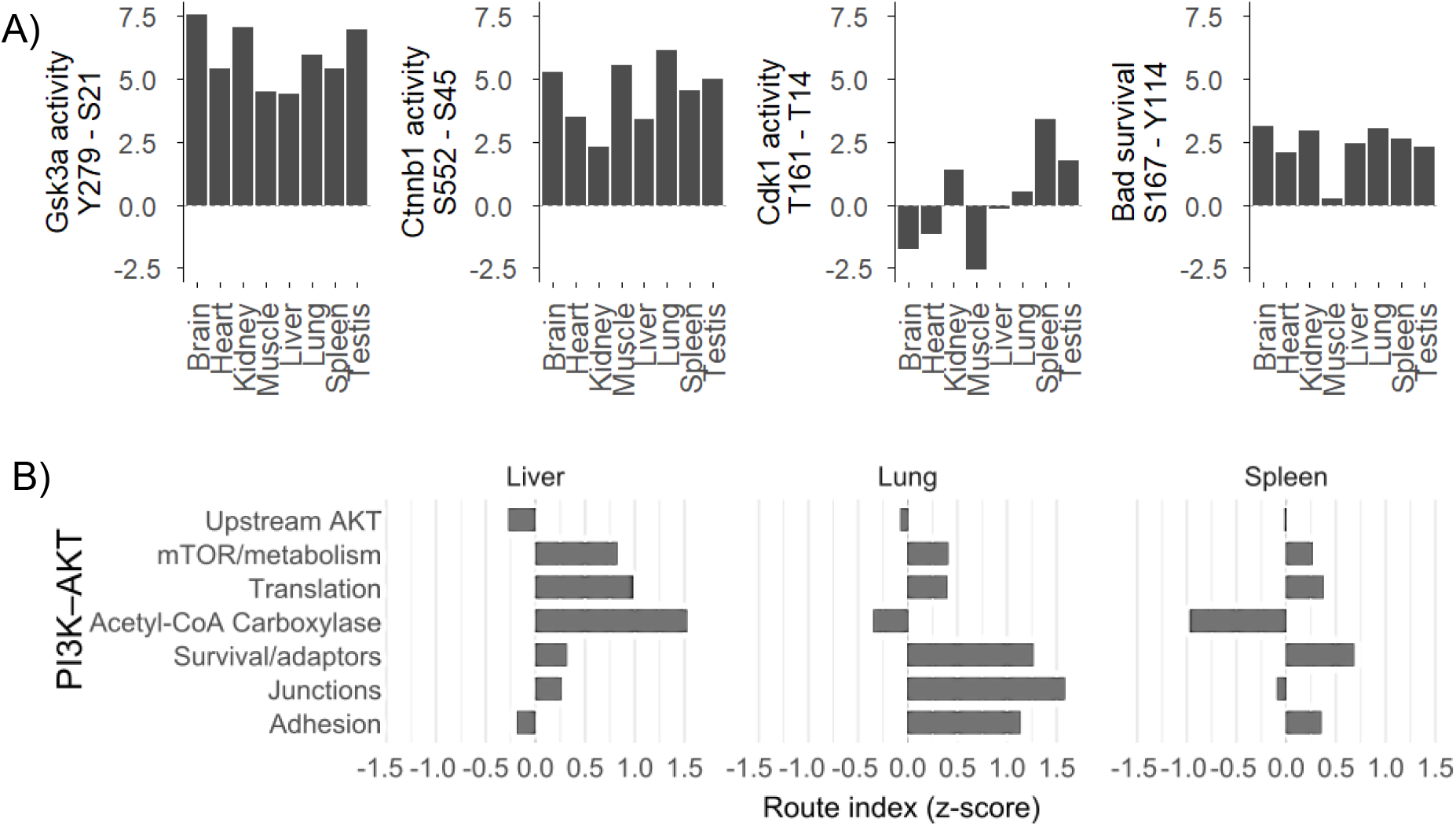
Directional kinase activity indices and PI3K-Akt pathway routing across organs. A) Directional activity indices for Gsk3a, Ctnnb1, Cdk1, alongside a Bad survival index, reflect the difference between the organ-level mean intensities of activating versus inhibitory phosphosites, calculated from the median-normalised, log2-transformed values. B) PI3K-Akt routing fingerprints for the liver, lung, and spleen were generated by averaging site-wise z-scores across predefined functional modules. Bar plots depict the calculated route index (z-score) per organ; positive and negative values represent above-average and below-average pathway activation, respectively, relative to the global cross-organ mean.

We next evaluated whether antagonistic indices report cell-cycle activation versus inhibition. For Cdk1, the activating T161 site was elevated in spleen and testis, whereas the inhibitory T14 site was prominent in testis and skeletal muscle (**Figure 1**). The T161–T14 index was highest in spleen and testis, and lowest in brain and skeletal muscle, consistent with proliferative versus post-mitotic tissue states (**Figure 2A**). Negative index values in brain and skeletal muscle indicate the dominance of inhibitory phosphorylation, consistent with Cdk1 being held inactive in these terminally differentiated tissues. FFPE-derived antagonistic indices thus resolve both activation and suppression of a core cell-cycle kinase and track directional shifts in its activity with tissue proliferation status.

To assess whether similar principles apply to survival signalling, we constructed a Bad survival index (S167-Y114). S167 phosphorylation is associated with pro-survival signalling, whereas Y114 is linked to pro-apoptotic regulation, and both sites were highest in the lung (**Figure 1**). The Bad survival index identified skeletal muscle as having a comparatively lower survival bias, consistent with reduced Bad-mediated survival signalling (**Figure 2A**). Antagonistic phosphosites can therefore be combined into directional indices that resolve the balance between pro-survival and pro-apoptotic signals in a tissue-resolved manner, extending this framework beyond kinases to survival regulators.

### Pi3k–Akt signalling is routed into distinct downstream branches across organs

We then asked whether FFPE phosphoproteomes capture how shared upstream signals are partitioned into distinct downstream branches in a tissue-specific manner. We defined branch-level “routing” indices for Pi3k-Akt signalling by averaging site-wise z-scores across curated effector modules, including Mtor/metabolism, translation, junctions, adhesion, and survival/adaptor signalling. Here, response strength corresponds to the mean *z*-scored phosphorylation within each functional branch, providing a measure of relative pathway usage across organs rather than absolute activity, allowing for descriptive ranking of pathway engagement.

These routing indices revealed clear tissue-specific allocation of Pi3k-Akt output. In the liver, signalling was preferentially directed into acetyl-CoA carboxylase, Mtor-linked metabolic control, and translation modules, consistent with the liver’s central role in lipogenesis and protein synthesis (**Figure 2B**). In the lung, the same upstream input was routed predominantly to junctional, adhesion, and survival/adaptor branches, compatible with the emphasis on epithelial-barrier integrity and pro-survival signalling in respiratory tissue. In the spleen, routing favoured survival/adaptor and adhesion modules, with comparatively limited engagement of the acetyl-CoA carboxylase branch, indicating a bias towards immune signalling and cell–cell interaction over lipid synthesis. FFPE phosphoproteomes thus capture differential allocation of signalling output across downstream branches of a shared pathway axis.

Isoform-resolved phosphosite measurements further supported branch-specific routing. Acaca S79 was enriched in the liver, whereas Acacb S219 was highest in the heart and skeletal muscle, consistent with cytosolic versus mitochondrial acetyl-CoA carboxylase usage in lipogenic versus oxidative tissues (**Figure 1**). These two phosphosites are distinguished by non-overlapping sequence windows, supporting isoform specific assignment at the phosphosite level. FFPE-derived data therefore resolve not only which branch receives preferential Pi3k–Akt input, but also which isoforms within that branch are engaged across organs

### FFPE phosphoproteomes delineate a kinase-by-organ activity landscape

To determine whether these routing differences are reflected at the level of upstream kinase activity, we constructed a proteome-normalised kinase-by-organ activity matrix. Using kinase-substrate relationship annotations established via potency-coherence analysis^52^ and proteome-normalised phosphosite intensities, we then aggregated phosphorylation levels across matched substrates for each kinase using median-centred values per sample (**Supplementary Table S5/6**). This shifts the analysis from individual phosphosites, which FFPE phosphoproteomes may not preserve uniformly, to composite, pathway-level readouts that treat each kinase as a functional unit rather than a collection of isolated sites. As with the antagonistic indices above, these composite scores are inferred kinase substrate activity scores derived from substrate phosphorylation and should be interpreted as rank-order comparisons of relative pathway engagement across organs rather than direct measurements of catalytic activity.

The resulting kinase-by-organ activity matrix revealed highly tissue-resolved signalling profiles across rat organs based on relative z-score enrichments (**Figure 3**). Growth, translational, and DNA-damage checkpoint kinases displayed pronounced tissue specificity. The testis showed coordinated enrichment of inferred Akt, Mtor, S6k, Sgk, and Rsk activity, alongside the checkpoint kinases Atm and Chek2 and the cell-cycle kinases Cdk1/2. This signature aligns with the high translational demands of continuous spermatogenesis and the DNA-damage surveillance required during meiotic recombination. Her-family and Src-family activity scores were also highest in the testis, consistent with growth-factor-driven regulation of spermatogenesis and the requirement for tight control of cell polarity, cytoskeletal organisation, and protein trafficking in Sertoli and germ cells. Conversely, the brain presented elevated inferred activity of Erk, Rsk, Akt, Mtor, and S6k, paired with structural and transcriptional regulators including Fak, ErbB-associated Her kinases, Aurkb, and Cdk9/12/13. This combination points to a signalling environment dominated by receptor-adhesion signalling at neuroglial contacts and sustained transcriptional elongation underlying activity-dependent gene expression.

**Figure 3.**
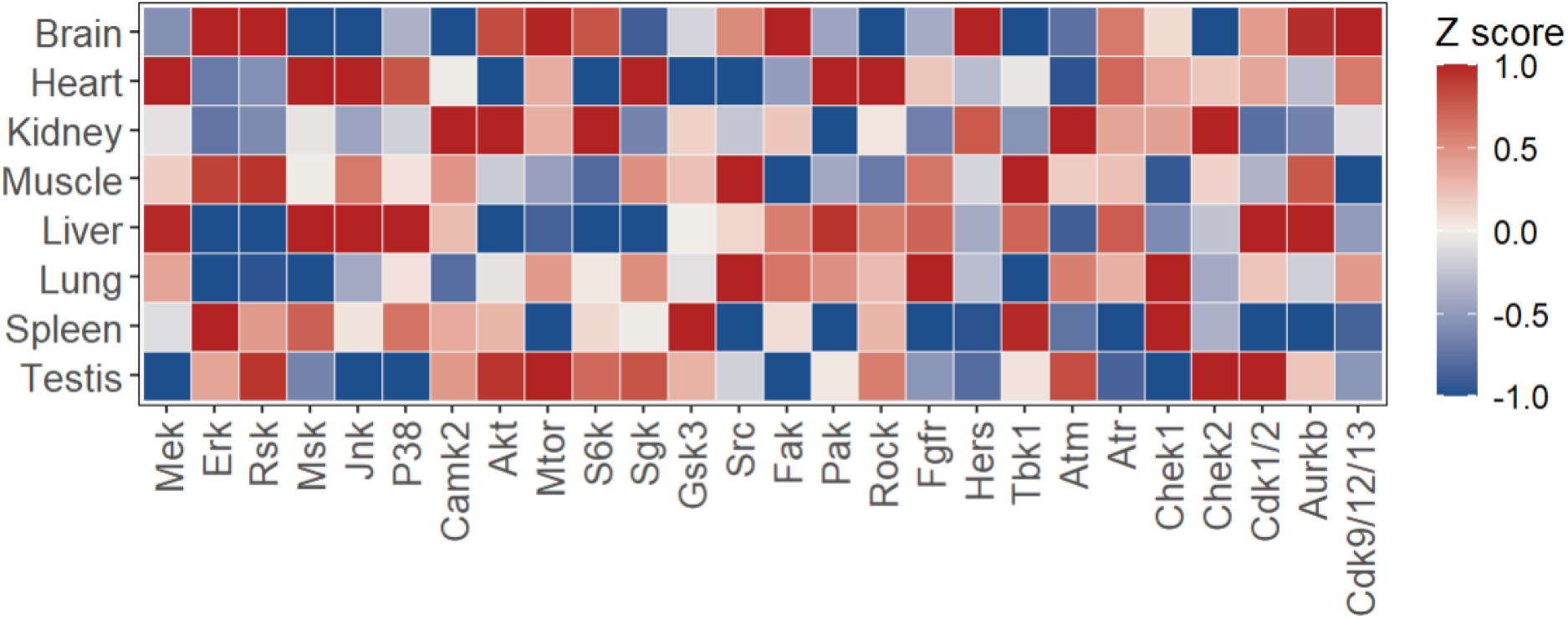
Proteome-normalised kinase-by-organ activity matrix across eight healthy rat organs. The matrix illustrates inferred kinase substrate activity scores, computed using curated kinase–substrate interactomes. To isolate genuine phosphorylation dynamics, site-level phosphosite intensities were strictly normalised to their respective parent protein abundances prior to scoring. The matrix displays organs as rows and kinase families as columns. For each kinase family, activity scores were first z-scored across the eight organs. Because structural protein dominance in certain tissues can introduce a shared, organ-wide technical baseline shared across kinase families, the median z-score across all kinase families was calculated for each organ and subtracted from every individual kinase family’s z-score in that organ, removing this shared baseline. The resulting offset-corrected values were then re-z-scored across organs for each kinase family, yielding the between-organ activity score displayed here (see Methods). Red and blue denote relative kinase activation and suppression, respectively. Z-scores have been clipped to ±1 to enhance visual clarity.

Stress-activated Mapk networks and cytoskeletal regulators demonstrated distinct compartmentalisation across the liver and heart. The liver displayed elevated inferred Mek, Msk, Jnk, p38, and Pak activity, which co-occurred with high inferred activity of Cdk1/2 and Aurkb. This profile reflects the liver’s dual capacity for stress-responsive detoxification and cell-cycle-competent regenerative turnover. Fgfr-associated activity was elevated in the heart, compatible with Fgf-mediated growth-factor signalling in cardiomyocytes. The heart similarly presented elevated inferred Mek, Msk, Jnk, p38, and Pak activity but coupled these with Sgk and Rock, indicating active cytoskeletal and contractile-apparatus remodelling. Notably, the heart lacked the Cdk1/2 and Aurkb activation seen in the liver, demonstrating that while these organs share a core stress-Mapk axis, the heart diverges downstream by coupling this axis exclusively to Rho-family cytoskeletal effectors rather than regenerative cell-cycle machinery.

Further organ-specific compartmentalisation was observed across metabolic and epithelial-surveillance networks. The kidney was enriched for inferred Camk2, Akt, S6k, Atm, and Chek2 activity, representing a functional coupling of calcium-dependent signalling, translational control, and genome-protective surveillance. In the spleen, inferred Erk and Gsk3 activity was highly elevated alongside the innate-immune kinase Tbk1 and Chek1, consistent with proliferative Mapk signalling, metabolic switching, and rapid lymphocyte turnover. Fgfr-associated activity was also elevated in the spleen, compatible with Fgf-mediated growth-factor signalling in immune cells. The lung displayed enrichment of inferred Src, Fgfr, and Chek1 activity, pointing to receptor tyrosine kinase-driven epithelial maintenance coupled with proliferative checkpoint control.

Structural and focal-adhesion pathways were distinctly partitioned between the two striated, contractile tissues. Skeletal muscle exhibited high inferred Erk and Rsk activity alongside the adhesion kinase Src, Tbk1, and Aurkb. This contrasted with the heart’s Pak/Rock-dominated cytoskeletal profile, and its reliance on the Mek–Msk–Jnk–p38 stress axis. These divergent signatures reveal that despite both being highly contractile tissues, skeletal muscle and heart engage substantially different upstream stress/growth modules (Erk/Rsk-driven in muscle vs. Jnk/p38-driven in heart) converging on distinct cytoskeletal effectors.

### FFPE and FF phosphoproteomes integrate into a shared coordinate space dominated by organ identity

We sought to determine whether FFPE and FF phosphoproteomes can be mapped into a shared phosphosite space in which variance is primarily driven by organ identity rather than preservation mode. We defined successful integration as the condition wherein organ identity accounts for the dominant fraction of total variance, with preservation mode contributing only a minor component. Under these conditions, FFPE and FF datasets can be treated as mutually compatible views of the same underlying signalling architecture, enabling FF models and clinical FFPE cohorts to be analysed within a unified map.

We combined FFPE phosphoproteomes from eight healthy Wistar rat organs with an external FF dataset from the same eight organs in Sprague–Dawley rats, yielding a joint matrix of 58,631 phosphosites mapped to 6,068 proteins and 5,787 genes (**Figure 4A, Supplementary Table S7**). Of these, 54,710 sites were detected in FFPE and 28,888 in FF, with 86% of FF sites also observed in FFPE based on exact site positional overlap. This roughly two-fold difference in total site counts reflects differences in acquisition depth and instrument sensitivity between the two studies, rather than a fundamental difference in the phosphoproteomes accessible from each preservation mode. The FFPE map presented here was generated with substantially greater analytical depth than the external FF dataset. This indicates that the vast majority of FF-detectable phosphosites are recoverable from FFPE tissue, and that both preservation modes sample essentially the same high-confidence phosphosite space, with divergence concentrated at the detection limit rather than reflecting distinct underlying biology. This overlap included organ-specific signatures in core cascade pathways, demonstrating that key regulatory axes are preserved across preservation methods. We note that this comparison spans not only preservation modes but also rat strains and independent cohorts; the dominance of organ identity over preservation-mode variance therefore represents a highly conservative estimate of FFPE fidelity.

**Figure 4.**
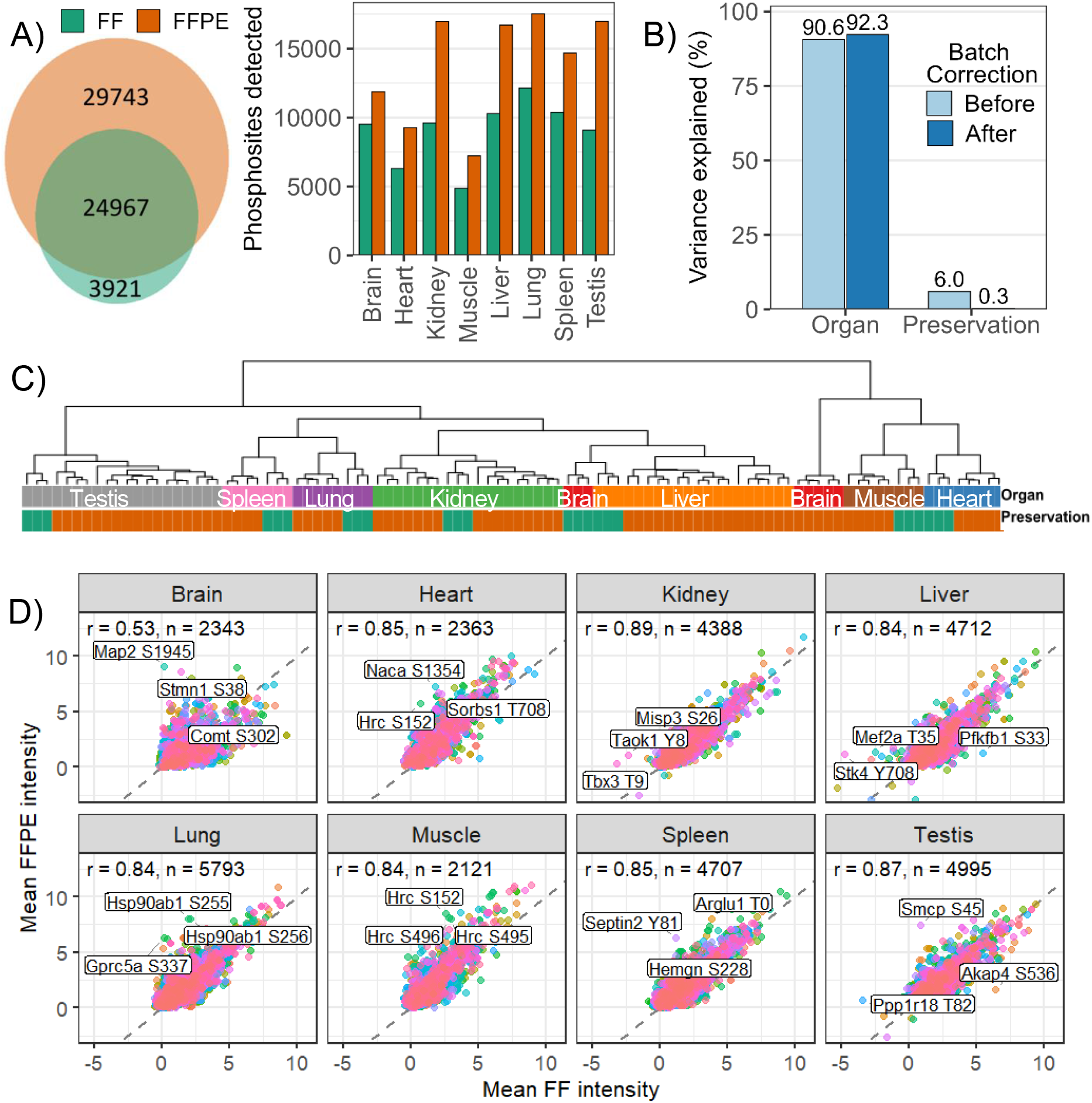
Integration of FFPE and FF phosphoproteomes into a unified, organ-resolved map. (A) Global intersection of phosphosites detected in FFPE and FF tissues, accompanied by per-organ identification counts across both preservation modes. (B) Principal variance component analysis quantifying the fraction of total dataset variance explained by organ identity versus preservation mode, both prior to and following ComBat batch correction. (C) Unsupervised hierarchical clustering of FFPE and FF samples post-batch correction with non-imputed values, demonstrating robust segregation by biological tissue rather than technical preparation. (D) Cross-preservation quantitative concordance. Per-organ scatter plots depict the Pearson correlation between mean log2 median-centred intensities in FFPE and FF samples, assessed exclusively across non-imputed phosphosites to ensure correlations reflect genuine biological conservation.

To examine preservation-specific signals, we classified phosphosites as shared (detected in both FF and FFPE), FF-unique, or FFPE-unique and compared their corrected intensities. Shared sites were more abundant (median abundance of 0.65 log2), whereas FF-unique and FFPE-unique sites exhibted lower medians (−0.79 and −0.77 log2, respectively), such that shared sites were ∼1.5 log2 units (∼2.8-fold) more intense than mode-specific sites (**Supplementary Figure S1**). Mode-specific sites were therefore enriched for low-abundance signals near the analytical detection limit, consistent with most such identifications arising from sensitivity differences and stochastic sampling of low-abundance phosphopeptides rather than from genuinely distinct phosphoproteomes in FF versus FFPE tissue.

To address whether computational batch correction might over-correct and mask genuine preservation-related signals, we applied principal variance component analysis to the matrices both before and after ComBat adjustment (**Supplementary Tables S8 and S9**). Notably, organ identity was already the overwhelmingly dominant driver of the dataset prior to batch correction, accounting for 90.6% of the total variance compared to only 6.0% for preservation mode (**Figure 4B/C, Supplementary Figure S2**). To evaluate the impact of this correction on FF versus FFPE concordance, we analysed the exact mathematical shifts applied to the data. The median absolute per-site correction was 0.0037 log2 units across phosphosites, indicating that ComBat applied minor adjustments relative to the biological signal range (**Supplementary Figure S3**). Because the biological signal overwhelmingly dominated the raw data, and the applied algorithmic adjustments were minimal, these results confirm that ComBat successfully mitigated minor technical matrix effects without over-correcting or erasing genuine biological signals. This indicates that the shared biological architecture between FF and FFPE is a native feature of the data rather than an artefact of normalisation. After batch correction, organ identity explained 92.3% of the variance, while preservation mode was reduced to 0.3%.

To rigorously quantify the reproducibility of these measurements without the confounding influence of data imputation, we first examined within-preservation Pearson correlations (r) between biological and technical replicates (**Supplementary Table S10**). Median correlations were high in both modes (FF r ≥ 0.92–0.95, FFPE r ≥ 0.91–0.96), indicating that variability introduced by sample preparation, enrichment, and LC–MS analysis is modest relative to the underlying biological signal. Next, we evaluated the per-organ cross-preservation Pearson correlation coefficients strictly using non-imputed phosphosites quantified in at least two biological replicates in both the FF and FFPE cohorts. While global variance decomposition requires a fully imputed matrix to compute principal components, masking all imputed values for this correlation analysis ensured our assessment reflected measured biological signals rather than algorithmic artefacts.

Across seven of the eight organs, global native phosphosite correlations were highly concordant (r = 0.84–0.89), with brain tissue as a notable exception (r = 0.53; **Figure 4D**). The comparatively lower brain concordance likely reflects the exceptional sensitivity of brain tissue to pre-analytical delay as it is known to undergo rapid post-mortem autolysis and enzymatic degradation compared with most other organs. This heighted sensitivity to ischemia duration and fixation kinetics may be compounded by the fact that synaptic phosphorylation networks are themselves highly dynamic and rapidly turned over. This represents a genuine biological and pre-analytical limitation of FFPE phosphoproteomics in neuronal tissue. Restricting the analysis exclusively to non-imputed phosphotyrosine sites yielded strong cross-preservation concordance across organs (r = 0.66 to 0.86; **Supplementary Figure S4**). While these native correlations are naturally lower than those derived from fully imputed matrices due to the removal of concordant lower-limit-of-detection values, they indicate that the quantified, high-confidence phosphosite landscape is biologically conserved between archival and FF processing.

Finally, we investigated the top 30 phosphosites exhibiting the largest quantitative divergence between FFPE and FF tissue after batch correction. Notably, no single phosphosite demonstrated a consistent, unidirectional abundance bias across all organs based on preservation mode (**Supplementary Figure S5, Supplementary Table S11**).

Taken together, the overlap analysis, abundance stratification, variance decomposition, clustering, within-mode reproducibility, and native per-site FF–FFPE correlations demonstrate that FFPE-derived phosphoproteomes integrate closely with FF datasets into a single, organ-resolved phosphosite map. These results provide independent evidence, across cohorts and rat strains, that organ identity rather than preservation mode governs phosphosite variation, supporting the use of FFPE-based studies of human disease within a reference frame originally defined in FF tissue.

## Discussion

The primary objective of this study was to determine whether FFPE tissue can support phosphosite-resolved signalling analyses with sufficient biological fidelity to serve as a map-scale resource. Our data demonstrate that FFPE phosphoproteomes predominantly recapitulate organ-specific signalling architectures rather than fixation artefacts, with one notable exception (brain) discussed below. By successfully integrating FFPE and FF datasets into a shared coordinate space, we support archival FFPE as a viable substrate for retrospective, systems-level signalling analysis, pending extension to human cohorts in the future.

FFPE yielded 54,710 phosphosites, substantially exceeding reported by FF studies^4, 13, 53, 54^ encompassing broad coverage across receptors, kinase cascades, and nuclear regulators. The resulting organ-resolved patterns aligned with known physiology, confirming that biological identity remains the primary driver of signalling architecture despite formaldehyde crosslinking. Furthermore, evaluating antagonistic phosphosite pairs on Gsk3a, Ctnnb1, Cdk1, and Bad permitted the calculation of activating-minus-inhibitory indices for pathway engagement, cell-cycle progression, and survival bias. Alongside composite PI3K-Akt routing indices and proteome-normalised kinase activity scores, these directional metrics confirm that FFPE material supports mechanistic, functional interpretations of signalling networks rather than basic pathway detection.

Integration with an independent, multi-organ FF dataset provided orthogonal validation of the FFPE-derived signalling architecture. Notably, organ-level variance remained dominant despite cohort and strain differences. While previous studies have documented the physicochemical biases induced by preanalytical factors and formaldehyde crosslinking^28, 55, 56^, our variance decomposition shows that, under controlled pre-analytical conditions, preservation-associated effects are computationally separable from biological variance. This establishes a framework to quantify not merely the presence of a pathway, but the specific engagement strength of key kinases across varied tissue contexts.

These findings establish the healthy-tissue baseline against which tumour-associated kinase rewiring could, in principle, be benchmarked. Unlike genomic alterations which denote pathway potential, phosphorylation reflects the real-time functional state of a tumour. Subject to validation in human FFPE tumour cohorts with defined pre-analytical parameters, the biological fidelity demonstrated here raises the prospect of retrospective signalling analysis in archived pathology material. Realising this potential requires a shift from single-site analyses to curated substrate signatures that infer kinase activity and network topology.^57–59^ Our kinase and pathway scores demonstrate the feasibility of this signature-based approach in FFPE tissue. Because such metrics aggregate signal across multiple substrates, they may be less sensitive to the loss or distortion of individual sites than single-epitope assays however this would need to be confirmed in matched tumour/adjacent-normal FFPE cohorts. As a proof of concept, these substrate-level kinase scores suggest a framework that, if validated in clinical cohorts, could support retrospective biomarker discovery in completed trial archives.

Methodologically, FFPE phosphoproteomics complements phospho-specific immunohistochemistry. While immunohistochemistry provides spatially resolved single-epitope markers, FFPE phosphoproteomics delivers multi-site, pathway-level signatures that quantify entire signalling axes in parallel. This methodology introduces a kinase-centric interpretation layer to histological collections, converting static tissue archives into maps of targetable signalling dependencies.

Several limitations outline necessary future work. First, the external FF comparator dataset differs from our FFPE workflow not only in preservation mode but also in animal strain, sample preparation, and LC-MS conditions. Because these factors vary jointly rather than independently, preservation mode is confounded with multiple other methodological differences between studies. This means the between-mode variance we report is a conservative combined upper bound on the effect of preservation alone. The true preservation-specific effect is therefore likely smaller than our reported estimate, meaning the concordance between preservation modes is, if anything, understated by this analysis. Second, the map is derived from rat models and these findings must be validated in human cohorts. Third, strict pre-analytical controls (e.g. cold ischemia and fixation times) were maintained here, which may not reflect the variable conditions of routine clinical pathology. Fourth, the current eight-organ baseline must be expanded to capture the complete spectrum of mammalian signalling. Fifth, brain tissue exhibited substantially lower cross-preservation concordance (r = 0.53 vs. r = 0.84–0.89 for other organs), likely reflecting the exceptional sensitivity of synaptic phosphorylation to pre-analytical delay. This warrants caution when applying FFPE phosphoproteomics to neural tissue studies. Lastly, this bulk-tissue approach sacrifices spatial resolution: phosphopeptide enrichment still requires substantial tissue input, precluding region- or cell-type-resolved analysis. Improving enrichment sensitivity and reduced input requirements make spatially resolved FFPE phosphoproteomics an increasingly realistic near-term goal.

A critical next step is translation of this framework to clinical human FFPE cohorts. This requires the generation of healthy-tissue phosphoproteome references across diverse biobanks, analogous to existing transcriptome and proteome maps^60–64^, to establish quantitative acceptance criteria and stable baselines. Such references are essential for preventing tissue-of-origin effects from confounding disease signals. Within this framework, FFPE-derived routing readouts could position patient tumours relative to healthy baselines, revealing pathological kinase rewiring and targetable vulnerabilities.

In conclusion, FFPE phosphoproteomics provides a foundation for organ-resolved signalling analysis in preclinical and translational settings. When pre-analytical variation and tissue-of-origin differences are carefully controlled, FFPE phosphoproteomics supports organ-resolved, systems-level signalling analysis, offering a route to mine existing clinical trial archives. Subject to validation in human cohorts, it could complement genomics and transcriptomics for mapping functional signalling states.

## Author contributions

EMH: Led the conceptualisation, methodology design, formal analysis, investigation, data curation, secured funding for the PhD exchange, wrote the original draft and led all subsequent revisions and editing. MS: Contributed to methodology, manuscript editing and review. NS: Manuscript editing and review. PGH: Supervision and access to resources. PJR: Supervision and access to resources. BK: Contributed to supervision, resources, manuscript editing and review.

## Acknowledgements

We thank all members of the Kuster lab for technical assistance and fruitful discussions. We thank Stephen Eckert and Johanna Tushaus for their expertise in FFPE tissue processing and mass spectrometry. We thank Magnus Huusfeldt, Joel Vej-Nielsen, Stoyan Stoychen, Dorte Bekker-Jensen and Nicolai Bache from Evosep and ReSyn Biosciences for providing EMH training on the Evosep One and some resources used in this study (stainless steel emitters and analytical columns).

This work backed by ProCan® was supported by the Australian Cancer Research Foundation, Cancer Institute New South Wales (NSW) (2017/TPG001, REG171150), NSW Ministry of Health (CMP-01), The University of Sydney, Cancer Council NSW (IG 18-01), Ian Potter Foundation, the Medical Research Futures Fund (MRFF-PD), National Health and Medical Research Council (NHMRC) of Australia European Union grant (GNT1170739, a companion grant to support the European Commission’s Horizon 2020 Program, H2020-SC1-DTH-2018-1, ’iPC-individualizedPaediatricCure’ [ref. 26121]), and National Breast Cancer Foundation (IIRS-18-164). The work was done under the auspices of a Memorandum of Understanding between Children’s Medical Research Institute and the U.S. National Cancer Institute’s International Cancer Proteogenomics Consortium (ICPC), that encourages cooperation among institutions and nations in proteogenomic cancer research in which datasets are made available to the public. E.M.H. is supported by an Australian Government Research Training Program (RTP) Scholarship, a Tour de Cure PhD Support Scholarship, and an Australian Graduate Women (AGW) Barbara Hale Fellowship. MS is supported by the Bavarian Cancer Research Center (BZKF) through the BZKF Lighthouse Omics, Genomics and Liquid Biopsy. NO is supported by the German Cancer Aid (Deutsche Krebshilfe), grant number 70116842. P.J.R. is supported by an NHMRC Fellowship (GNT1137064).

## Supplementary Material

### Supplementary Tables

Table S1: Sample annotations and MS files

Table S2: Phosphosite raw output containing 90,255 phosphosites on 6,079 protein groups. SwissProt and TrEMBL entries are listed separately

Table S3: FFPE phosphoproteome map. Phosphosite intensities are log2-transformed only; no normalisation, imputation, batch correction, or filtering applied

Table S4: FFPE phosphoproteome panel of signalling sites spanning major pathways across eight organs. Phosphosite intensities are median-normalised, log2-transformed, imputed, and z-scored

Table S5: FFPE proteome background. Protein group log2-transformed intensities per organ (no normalisation or imputation applied)

Table S6: FFPE proteome-normalised phosphosite intensities (log2 phospho − log2 proteome)

Table S7: Integrated FFPE and FF matrix. Phosphosite intensities are log2-transformed only; no normalisation, imputation, batch correction, or filtering applied

Table S8: FFPE and FF phosphoproteome map with organ-level mean values. Phosphosite intensities are median-normalised and log2-transformed only, with no imputation or batch correction; a minimum 10% detection threshold was applied.

Table S9: FFPE and FF phosphoproteome map with organ-level mean values. Phosphosite intensities are median-normalised, log2-transformed only, imputation removed after batch correction; a minimum 10% detection threshold was applied.

Table S10: Within-preservation Pearson correlations between biological and technical replicates, calculated on natively detected phosphosite intensities (imputed values excluded).

Table S11: Top 30 phosphosites exhibiting the largest quantitative divergence between FFPE and FF tissue after batch correction.

---

**Supplementary Figure S1.**
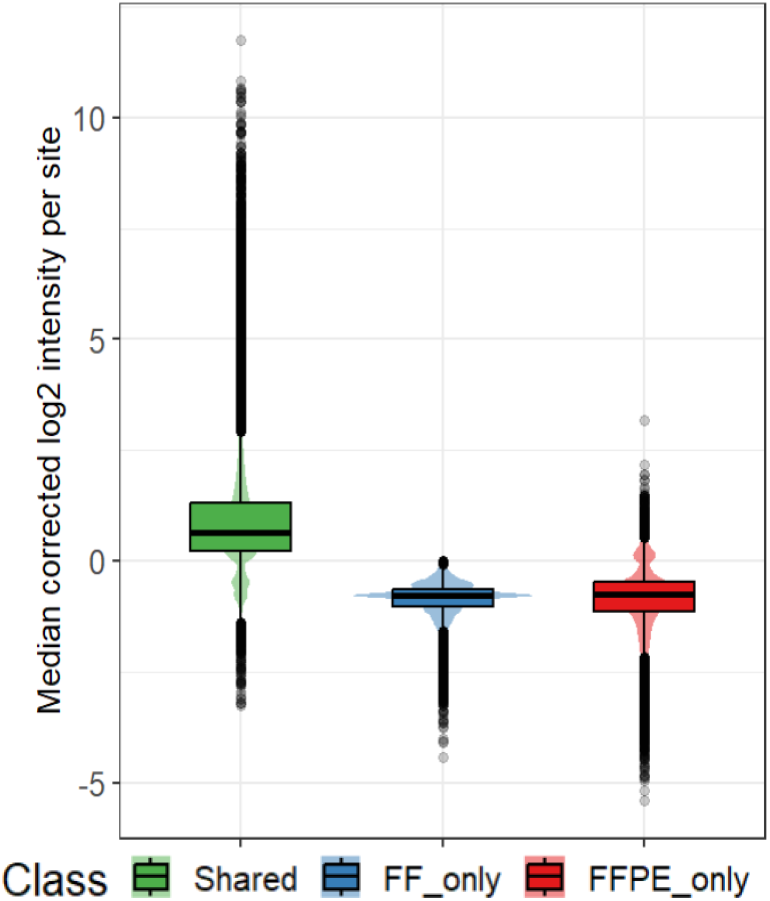
Abundance distributions of shared and preservation-specific phosphosites. Global intensity distributions are shown for phosphosites detected in both fresh frozen (FF) and formalin-fixed paraffin-embedded (FFPE) tissues, as well as sites unique to either FF or FFPE modes. Data represent fully processed intensities (median-normalised, log2-transformed, imputed, and batch-corrected) to illustrate the enrichment of mode-specific identifications near the analytical detection limit compared to highly abundant shared sites.

**Supplementary Figure S2.**
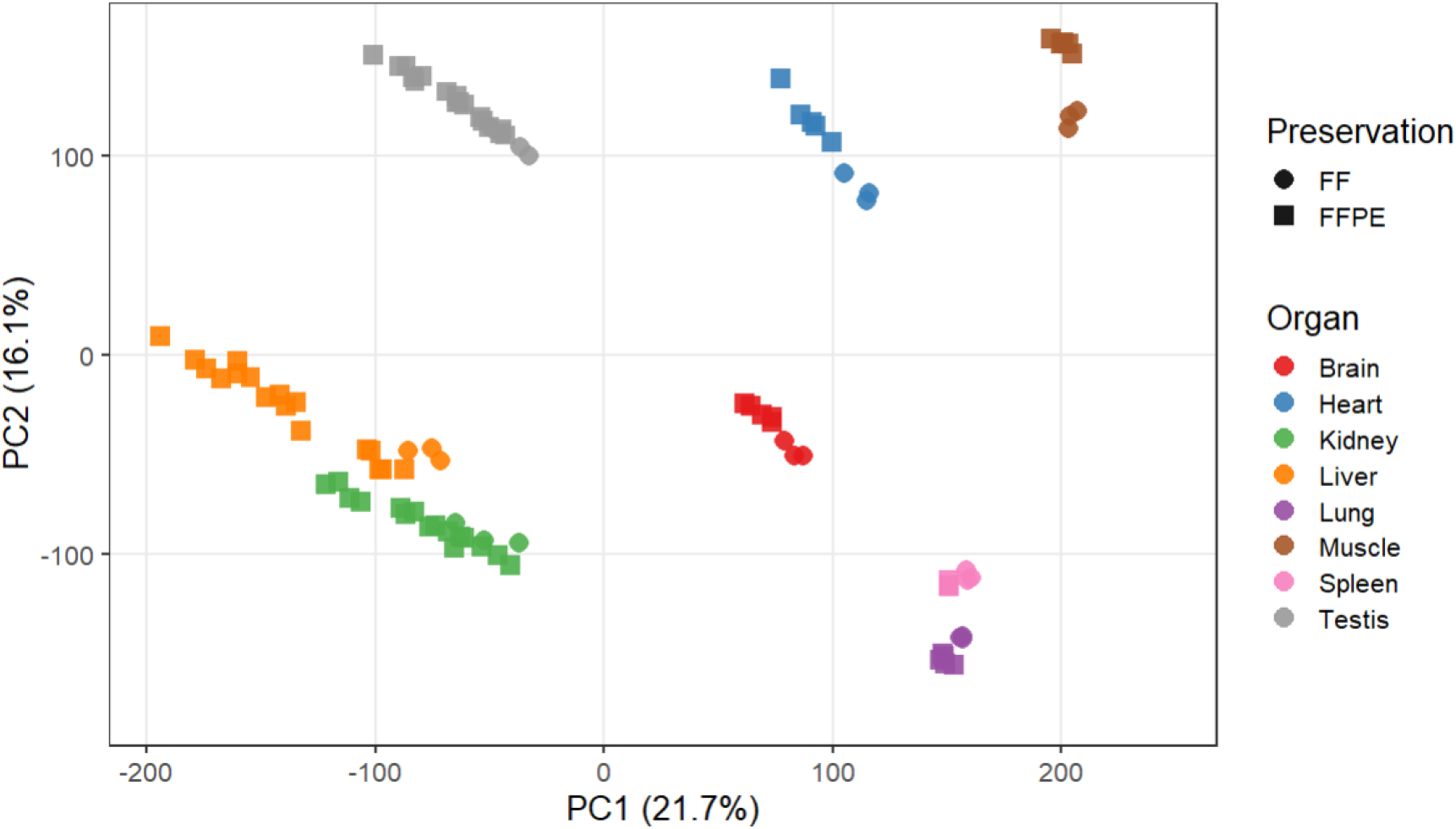
Principal component analysis (PCA) of the uncorrected phosphoproteome. PCA demonstrates the global variance and clustering of samples by rat organ and preservation mode prior to batch correction. The analysis is based on the median-normalised, log2-transformed, and imputed matrix containing 31,169 phosphosites, revealing that organ identity dominates dataset variance prior to algorithmic alignment.

**Supplementary Figure S3.**
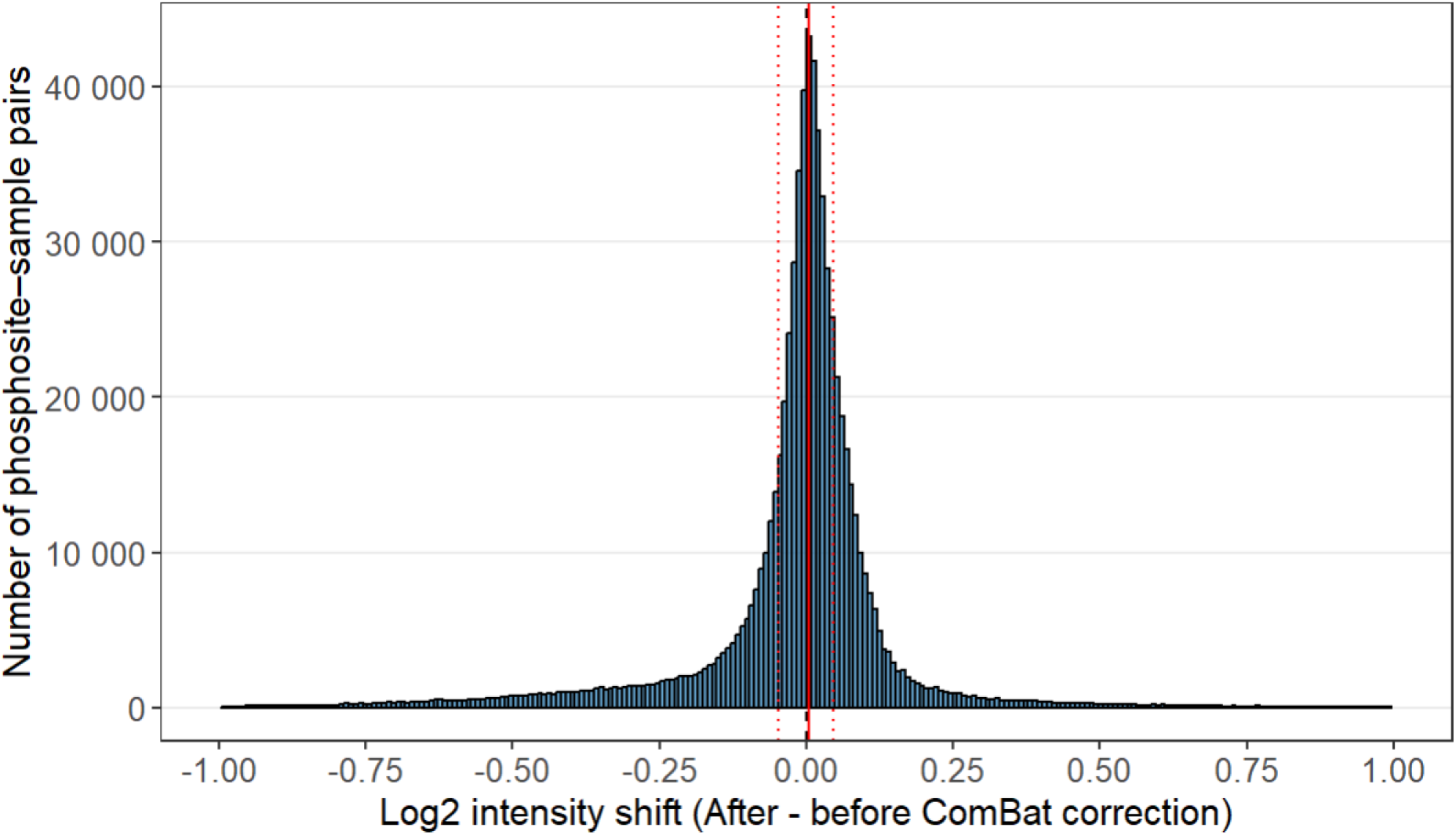
Magnitude of batch correction adjustments across the phosphoproteome. The distribution illustrates the median absolute shift applied to phosphosite intensities during ComBat batch correction across all organs. The solid red line designates the global median shift (0.0037 log2 units), and the dotted lines indicate the interquartile range (Q1 = −0.047, Q3 = 0.045), highlighting the minimal algorithmic intervention required to align the FF and FFPE datasets.

**Supplementary Figure S4.**
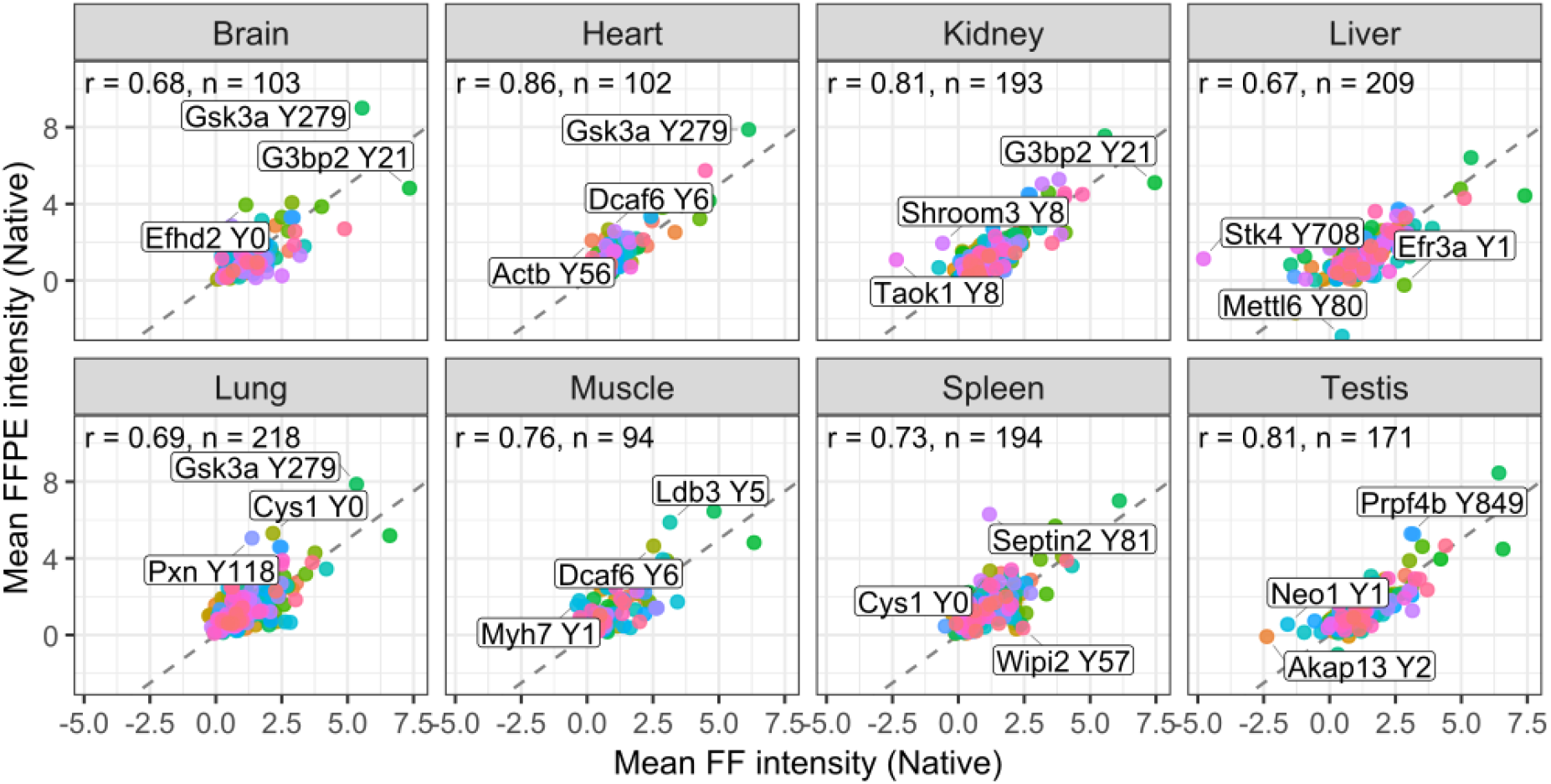
Cross-preservation concordance of the phosphotyrosine landscape. Per-organ scatter plots and Pearson correlation analyses compare the mean log2 median-centred, and batch corrected intensities between FFPE and FF tissues. To prevent algorithmic artefacts, this correlation evaluates only non-imputed quantified phosphotyrosine sites (excluding all imputed values), demonstrating the robust biological conservation of tyrosine regulatory networks across archival and fresh processing.

**Supplementary Figure S5.**
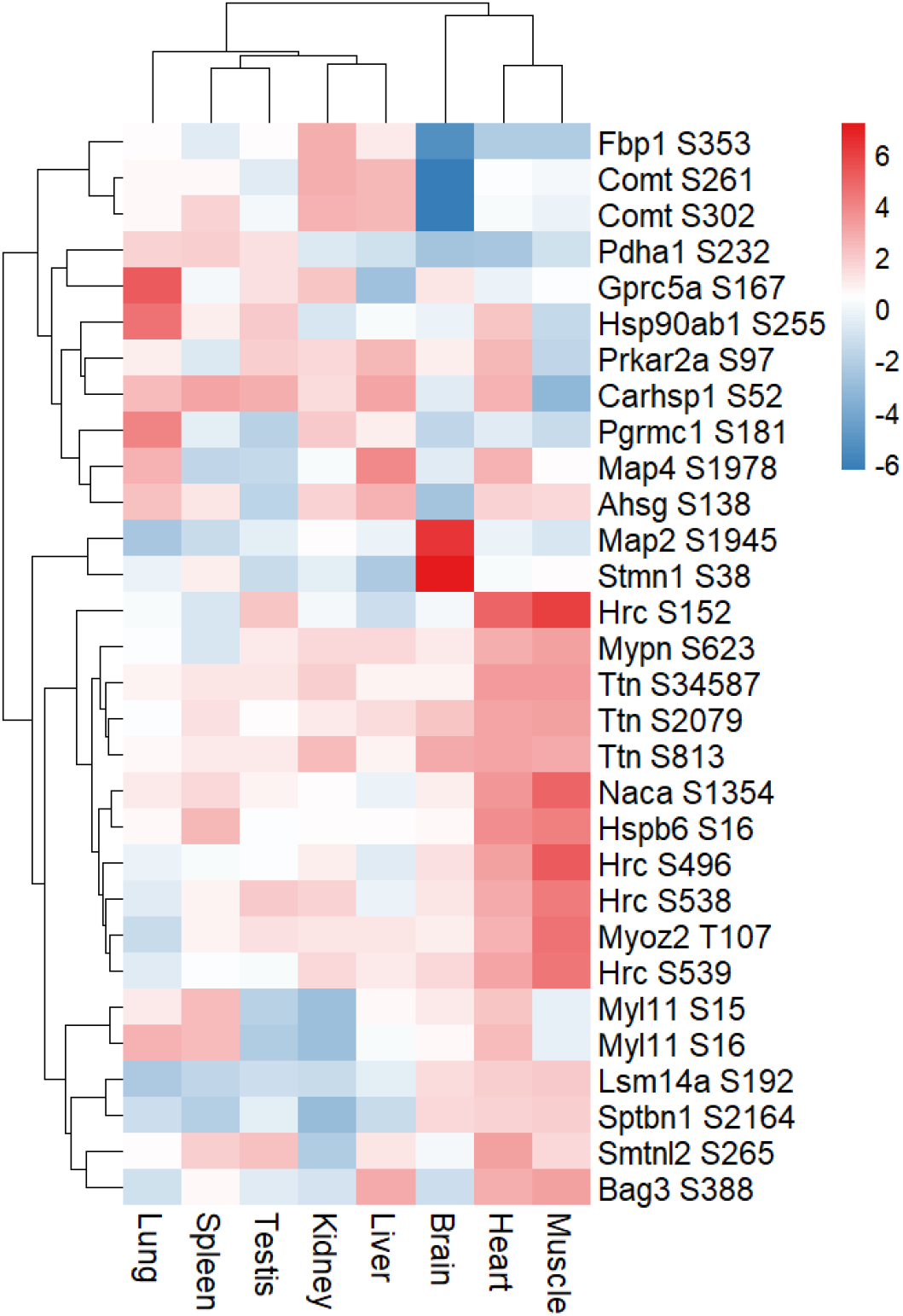
Top 30 phosphosites exhibiting the greatest quantitative divergence between FF and FFPE tissues. The data represents the phosphosites with the largest mean intensity differences (delta) between preservation modes following batch correction. The bidirectional distribution of these deltas confirms the absence of a consistent, unidirectional abundance bias driven by preservation mode across the organs.

## References

(1) Riley, N. M.; Coon, J. J. Phosphoproteomics in the Age of Rapid and Deep Proteome Profiling. Anal. Chem 2016, 88 (1), 74–94. DOI: 10.1021/acs.analchem.5b04123.

(2) Higgins, L.; Gerdes, H.; Cutillas, P. R. Principles of phosphoproteomics and applications in cancer research. Biochem. J 2023, 480 (6), 403–420. DOI: 10.1042/BCJ20220220.

(3) Koh, J. M. S.; Sykes, E. K.; Rukhaya, J.; Anees, A.; Zhong, Q.; Jackson, C.; Panizza, B. J.; Reddel, R. R.; Balleine, R. L.; Hains, P. G.;, et al. The effect of storage time and temperature on the proteomic analysis of FFPE tissue sections. Clin Proteom 2025, 22 (5). DOI: 10.1186/s12014-025-09529-5.

(4) Haines, M.; Thorup, J. R.; Gohsman, S.; Ctortecka, C.; Newton, C.; Rohrer, D. C.; Hostetter, G.; Mani, D. R.; Gillette, M. A.; Satpathy, S.;, et al. High-Throughput Proteomic and Phosphoproteomic Analysis of Formalin-Fixed Paraffin-Embedded Tissues. Mol Cell Proteomics 2025, 24 (9). DOI: DOI: 10.1016/j.mcpro.2025.101044.

(5) Xavier, D.; Lucas, N.; Williams, S. G.; Koh, J. M. S.; Ashman, K.; Loudon, C.; Reddel, R.; Hains, P. G.; Robinson, P. J. Heat ‘n Beat: A universal high-throughput end-to-end proteomics sample processing platform in under an hour. Anal. Chem 2024, 96 (27), 4093–4102. DOI: 10.1021/acs.analchem.3c04708.

(6) Humphries, E. M.; Loudon, C.; Hains, P. G.; Robinson, P. J. Quantitative comparison of deparaffinisation, rehydration and extraction methods for FFPE tissue proteomics and phosphoproteomics. Anal. Chem 2024, 96 (33), 13358–13370. DOI: 10.1021/acs.analchem.3c04479.

(7) Makhmut, A.; Qin, D.; Hartlmayr, D.; Seth, A.; Coscia, F. An Automated and Fast Sample Preparation Workflow for Laser Microdissection Guided Ultrasensitive Proteomics. Mol. Cell. Proteom 2024, 23 (5), 100750. DOI: 10.1016/j.mcpro.2024.100750.

(8) Tüshaus, J.; Eckert, S.; Fraefel, M.; Zhou, Y.; Pfeiffer, P.; Halves, C.; Fusco, F.; Weigl, J.; Hönikl, L.; Butenschön, V.;, et al. Towards routine proteome profiling of FFPE tissue: Insights from a 1,200 case pan-cancer study. EMBO. J 2024, 1–26–26. DOI: 10.1038/s44318-024-00289-w.

(9) Bader, J. M.; Deigendesch, N.; Misch, M.; Mann, M.; Koch, A.; Meissner, F. Proteomics separates adult-type diffuse high-grade gliomas in metabolic subgroups independent of 1p/19q codeletion and across IDH mutational status. Cell. Rep. Med 2023, 4 (1), 100877. DOI: 10.1016/j.xcrm.2022.100877.

(10) Mitsa, G.; Richard, V. R.; Majedi, Y.; Lafleur, J.; Aguilar-Mahecha, A.; Basik, M.; Borchers, C. H. Evaluation of a ’plug and play’ nanoflow liquid chromatography system for MS-based proteomic characterization of clinical FFPE specimens. Expert. Rev. Proteom 2023, 1–6. DOI: 10.1080/14789450.2023.2219844 From NLM Publisher.

(11) Tüshaus, J.; Sakhteman, A.; Lechner, S.; The, M.; Mucha, E.; Krisp, C.; Schlegel, J.; Delbridge, C.; Kuster, B. A region-resolved proteomic map of the human brain enabled by high-throughput proteomics. EMBO. J 2023, 42 (23), e114665. DOI: 10.15252/embj.2023114665.

(12) Kohale, I. N.; Burgenske, D. M.; Mladek, A. C.; Bakken, K. K.; Kuang, J.; Boughey, J. C.; Wang, L.; Carter, J. M.; Haura, E. B.; Goetz, M. P.;, et al. Quantitative analysis of tyrosine phosphorylation from FFPE tissues reveals patient specific signaling networks. Cancer Res 2021, 81 (14), 3930–3941. DOI: 10.1158/0008-5472.CAN-21-0214.

(13) Friedrich, C.; Schallenberg, S.; Kirchner, M.; Ziehm, M.; Niquet, S.; Haji, M.; Beier, C.; Neudecker, J.; Klauschen, F.; Mertins, P. Comprehensive micro-scaled proteome and phosphoproteome characterization of archived retrospective cancer repositories. Nat. Commun 2021, 12 (1), 3576. DOI: 10.1038/s41467-021-23855-w.

(14) Eckert, S.; Chang, Y. C.; Bayer, F. P.; The, M.; Kuhn, P. H.; Weichert, W.; Kuster, B. Evaluation of Disposable Trap Column nanoLC-FAIMS-MS/MS for the Proteomic Analysis of FFPE Tissue. J. Proteome. Res 2021, 20 (12), 5402–5411. DOI: 10.1021/acs.jproteome.1c00695.

(15) Mantsiou, A.; Makridakis, M.; Fasoulakis, K.; Katafigiotis, I.; Constantinides, C. A.; Zoidakis, J.; Roubelakis, M. G.; Vlahou, A.; Lygirou, V. Proteomics Analysis of Formalin Fixed Paraffin Embedded Tissues in the Investigation of Prostate Cancer. J. Proteome. Res 2020, 19 (7), 2631–2642. DOI: 10.1021/acs.jproteome.9b00587.

(16) Oberhuber, M.; Pecoraro, M.; Rusz, M.; Oberhuber, G.; Wieselberg, M.; Haslinger, P.; Gurnhofer, E.; Schlederer, M.; Limberger, T.; Lagger, S.;, et al. STAT3-dependent analysis reveals PDK4 as independent predictor of recurrence in prostate cancer. Mol. Syst. Biol 2020, (4), e9247. DOI: DOI: 10.15252/msb.20199247.

(17) Coscia, F.; Doll, S.; Bech, J. M.; Schweizer, L.; Mund, A.; Lengyel, E.; Lindebjerg, J.; Madsen, G. I.; Moreira, J. M. A.; Mann, M. A streamlined mass spectrometry-based proteomics workflow for large-scale FFPE tissue analysis. J. Pathol 2020, 251 (1), 100–112. DOI: 10.1002/path.5420.

(18) Kuras, M.; Woldmar, N.; Kim, Y.; Hefner, M.; Malm, J.; Moldvay, J.; Dome, B.; Fillinger, J.; Pizzatti, L.; Gil, J.;, et al. Proteomic Workflows for High-Quality Quantitative Proteome and Post-Translational Modification Analysis of Clinically Relevant Samples from Formalin-Fixed Paraffin-Embedded Archives. J. Proteome. Res 2020, 20 (1), 1027–1039. DOI: 10.1021/acs.jproteome.0c00850.

(19) Ostasiewicz, P.; Wisniewski, J. R. Chapter 10 A Protocol for Large-Scale Proteomic Analysis of Microdissected Formalin Fixed and Paraffin Embedded Tissue. In Methods Enzymol, 2017/01/23 ed.; Vol. 585; 2017; pp 159–176.

(20) Hughes, C. S.; McConechy, M. K.; Cochrane, D. R.; Nazeran, T.; Karnezis, A. N.; Huntsman, D. G.; Morin, G. B. Quantitative Profiling of Single Formalin Fixed Tumour Sections: proteomics for translational research. Sci. Rep 2016, 6, 34949. DOI: 10.1038/srep34949.

(21) Gustafsson, O. J.; Arentz, G.; Hoffmann, P. Proteomic developments in the analysis of formalin-fixed tissue. Biochim. Biophys. Acta 2015, 1854 (6), 559–580. DOI: 10.1016/j.bbapap.2014.10.003.

(22) Wakabayashi, M.; Yoshihara, H.; Masuda, T.; Tsukahara, M.; Sugiyama, N.; Ishihama, Y. Phosphoproteome Analysis of Formalin-Fixed and Paraffin-Embedded Tissue Sections Mounted on Microscope Slides. J. Proteome. Res 2014, 13 (2), 915–924. DOI: 10.1021/pr400960r.

(23) Fowler, C. B.; O’Leary, T. J.; Mason, J. T. Improving the Proteomic Analysis of Archival Tissue by Using Pressure-Assisted Protein Extraction: A Mechanistic Approach. J. Proteomics. Bioinform 2014, 7 (6), 151–157. DOI: 10.4172/jpb.1000315.

(24) Gámez-Pozo, A.; Sánchez-Navarro, I.; Ibarz Ferrer, N.; García Martínez, F.; Ashman, K.; Fresno Vara, J. Á. High-Throughput Phosphoproteomics from Formalin-Fixed, Paraffin-Embedded Tissues. Curr. Protoc. Chem. Biol 2012, 4 (2), 161–175. DOI: 10.1002/9780470559277.ch110242.

(25) Wiśniewski, J. R.; Ostasiewicz, P.; Duś, K.; Zielińska, D. F.; Gnad, F.; Mann, M. Extensive quantitative remodeling of the proteome between normal colon tissue and adenocarcinoma. Mol. Syst. Biol 2012, 8 (1), 611. DOI: 10.1038/msb.2012.44 (accessed 2024/11/20).

(26) Wolff, C.; Schott, C.; Porschewski, P.; Reischauer, B.; Becker, K. F. Successful protein extraction from over-fixed and long-term stored formalin-fixed tissues. Plos One 2011, 6 (1), e16353. DOI: 10.1371/journal.pone.0016353.

(27) Gamez-Pozo, A.; Sanchez-Navarro, I.; Calvo, E.; Diaz, E.; Miguel-Martin, M.; Lopez, R.; Agullo, T.; Camafeita, E.; Espinosa, E.; Lopez, J. A.;, et al. Protein phosphorylation analysis in archival clinical cancer samples by shotgun and targeted proteomics approaches. Mol. Biosyst 2011, 7 (8), 2368–2374. DOI: 10.1039/c1mb05113j.

(28) Ostasiewicz, P.; Zielinska, D. F.; Mann, M.; Wisniewski, J. R. Proteome, Phosphoproteome, and N-Glycoproteome Are Quantitatively Preserved in Formalin-Fixed Paraffin-Embedded Tissue and Analyzable by High-Resolution Mass Spectrometry. J. Proteome. Res 2010, 9 (7), 3688–3700. DOI: 10.1021/pr100234w.

(29) Becker, K.-F.; Schott, C.; Becker, I.; Höfler, H. Guided protein extraction from formalin-fixed tissues for quantitative multiplex analysis avoids detrimental effects of histological stains. Proteome. Clin. Appl 2008, 2 (5), 737–743. DOI: 10.1002/prca.200780106 (accessed 2024/11/19).

(30) Guo, T.; Wang, W.; Rudnick, P. A.; Song, T.; Li, J.; Zhuang, Z.; Weil, R. J.; DeVoe, D. L.; Lee, C. S.; Balgley, B. M. Proteome Analysis of Microdissected Formalin-fixed and Paraffin-embedded Tissue Specimens. J. Histochem. Cytochem 2007, 55 (7), 763–772. DOI: 10.1369/jhc.7A7177.2007.

(31) Hood, B. L.; Conrads, T. P.; Veenstra, T. D. Mass spectrometric analysis of formalin-fixed paraffin-embedded tissue: Unlocking the proteome within. Proteomics 2006, 6 (14), 4106–4114. DOI: 10.1002/pmic.200600016 (accessed 2024/11/10).

(32) Humphries, E. M.; Hains, P. G.; Robinson, P. J. Overlap of matched fresh-frozen and formalin-fixed, paraffin-embedded tissues for proteomics and phosphoproteomics. ACS Omega 2025, 10 (7), 6891–6900. DOI: 10.1021/acsomega.4c09289.

(33) Fröhlich, K.; Fahrner, M.; Brombacher, E.; Seredynska, A.; Maldacker, M.; Kreutz, C.; Schmidt, A.; Schilling, O. Data-Independent Acquisition: A Milestone and Prospect in Clinical Mass Spectrometry Based Proteomics. Mol Cell Proteomics 2024, 23 (8). DOI: 10.1016/j.mcpro.2024.100800 (accessed 2026/07/02).

(34) Mandell, J. W. Immunohistochemical assessment of protein phosphorylation state: the dream and the reality. Histochem. Cell. Biol 2008, 130 (3), 465–471. DOI: 10.1007/s00418-008-0474-z.

(35) Mandell, J. W. Phosphorylation State-Specific Antibodies: Applications in Investigative and Diagnostic Pathology. The American Journal of Pathology 2003, 163 (5), 1687–1698. DOI: 10.1016/S0002-9440(10)63525-0.

(36) George, B.; Haque, A.; Sahu, V.; Joldoshova, A.; Singh, Y.; Quinones, J. E.; George, S. K.; Amin, H. M. Enhancing Antigen Retrieval to Unmask Signaling Phosphoproteins in Formalin-fixed Archival Tissues. Appl. Immunohistochem. Mol. Morphol 2022, 30 (5). DOI: DOI: 10.1097/PAI.0000000000001022.

(37) Wang, N.; Zhang, L.; Ying, Q.; Song, Z.; Lu, A.; Treumann, A.; Liu, Z.; Sun, T.; Ding, Z. A reverse phase protein array based phospho-antibody characterization approach and its applicability for clinical derived tissue specimens. Sci. Rep 2022, 12 (1), 22373. DOI: DOI 10.1038/s41598-022-26715-9.

(38) Burns, J. A.; Li, Y.; Cheney, C. A.; Ou, Y.; Franlin-Pfeifer, L. L.; Kuklin, N.; Zhang, Z.-Q. Choice of Fixative Is Crucial to Successful Immunohistochemical Detection of Phosphoproteins in Paraffin-embedded Tumor Tissues. Journal of Histochemistry & Cytochemistry 2009, 57 (3), 257–264. DOI: DOI: 10.1369/jhc.2008.952911.

(39) Wolf, C.; Jarutat, T.; Vega Harring, S.; Haupt, K.; Babitzki, G.; Bader, S.; David, K.; Juhl, H.; Arbogast, S. Determination of phosphorylated proteins in tissue specimens requires high-quality samples collected under stringent conditions. Histopathology 2014, 64 (3), 431–444. DOI: DOI: 10.1111/his.12268.

(40) Mueller, C.; Edmiston, K. H.; Carpenter, C.; Gaffney, E.; Ryan, C.; Ward, R.; White, S.; Memeo, L.; Colarossi, C.; Petricoin, E. F., 3rd; et al. One-step preservation of phosphoproteins and tissue morphology at room temperature for diagnostic and research specimens. Plos One 2011, 6 (8), e23780. DOI: 10.1371/journal.pone.0023780.

(41) Vassilakopoulou, M.; Parisi, F.; Siddiqui, S.; England, A. M.; Zarella, E. R.; Anagnostou, V.; Kluger, Y.; Hicks, D. G.; Rimm, D. L.; Neumeister, V. M. Preanalytical variables and phosphoepitope expression in FFPE tissue: quantitative epitope assessment after variable cold ischemic time. Lab. Invest 2015, 95 (3), 334–341. DOI: 10.1038/labinvest.2014.139.

(42) Neumeister, V. M.; Anagnostou, V.; Siddiqui, S.; England, A. M.; Zarrella, E. R.; Vassilakopoulou, M.; Parisi, F.; Kluger, Y.; Hicks, D. G.; Rimm, D. L. Quantitative Assessment of Effect of Preanalytic Cold Ischemic Time on Protein Expression in Breast Cancer Tissues. J. Natl. Cancer. Inst 2012, 104 (23), 1815–1824. DOI: 10.1093/jnci/djs438 (accessed 11/17/2024).

(43) Neumeister, V. M.; Parisi, F.; England, A. M.; Siddiqui, S.; Anagnostou, V.; Zarrella, E.; Vassilakopolou, M.; Bai, Y.; Saylor, S.; Sapino, A.;, et al. A tissue quality index: an intrinsic control for measurement of effects of preanalytical variables on FFPE tissue. Lab Invest 2014, 94 (4), 467–474. DOI: 10.1038/labinvest.2014.7.

(44) Schneider, A.; Wortmann, J.; Jensen, C. B.; Duenas, L. E.; Teleanu, M.-V.; Sakhteman, A.; Hamood, F.; Bayer, F. P.; Stange, C.; Santoso, J. B.; et al. Prospective pan-cancer phosphoproteomics at clinical scale extends therapeutic options in precision oncology. bioRxiv 2026, 2026.2007.2008.737171. DOI: 10.64898/2026.07.08.737171.

(45) Jensen, C. B.; Sakhteman, A.; Hamood, F.; Schneider, A.; Woortman, J.; Teleanu, M.-V.; Horak, P.; Bayer, F. P.; Stange, C.; Huellein, J.; et al. TOPAS: phosphoproteome data analysis and decision support platform for molecular tumor boards. bioRxiv 2026, 2026.2007.2008.737143. DOI: 10.64898/2026.07.08.737143.

(46) Muller, T.; Kalxdorf, M.; Longuespee, R.; Kazdal, D. N.; Stenzinger, A.; Krijgsveld, J. Automated sample preparation with SP3 for low-input clinical proteomics. Mol. Syst. Biol 2020, 16 (1), e9111. DOI: 10.15252/msb.20199111.

(47) Humphries, E. M.; Xavier, D.; Ashman, K.; Hains, P. G.; Robinson, P. J. High-Throughput Proteomics and Phosphoproteomics of Rat Tissues Using Microflow Zeno SWATH. J. Proteome. Res 2024. DOI: 10.1021/acs.jproteome.4c00010.

(48) Bekker-Jensen, D. B.; Martinez-Val, A.; Steigerwald, S.; Ruther, P.; Fort, K. L.; Arrey, T. N.; Harder, A.; Makarov, A.; Olsen, J. V. A Compact Quadrupole-Orbitrap Mass Spectrometer with FAIMS Interface Improves Proteome Coverage in Short LC Gradients. Mol. Cell. Proteom 2020, 19 (4), 716–729. DOI: 10.1074/mcp.TIR119.001906.

(49) Bruderer, R.; Bernhardt, O. M.; Gandhi, T.; Miladinovic, S. M.; Cheng, L. Y.; Messner, S.; Ehrenberger, T.; Zanotelli, V.; Butscheid, Y.; Escher, C.;, et al. Extending the limits of quantitative proteome profiling with data-independent acquisition and application to acetaminophen-treated three-dimensional liver microtissues. Mol. Cell. Proteom 2015, 14 (5), 1400–1410. DOI: 10.1074/mcp.M114.044305.

(50) Consortium, U. UniProt: the Universal Protein Knowledgebase in 2023. Nucleic Acids Res 2023, 51, D523–D531. DOI: 10.1093/nar/gkac1052.

(51) L’Ecuyer, P.; Simard, R.; Chen, E. J.; Kelton, W. D. An Object-Oriented Random-Number Package with Many Long Streams and Substreams. Operations Research 2002, 50 (6), 1073–1075. DOI: 10.1287/opre.50.6.1073.358.

(52) Bayer, F. P.; Müller, J.; Kabella, N.; Abele, M.; Chang, Y.-C.; Estrada Duenas, L. D.; Jensen, C.; Hamood, F.; Lee, C.-Y.; Schneider, A.; et al. Chemical proteomics decrypts the kinases that shape the dynamic human phosphoproteome. bioRxiv 2025, 2025.2011.2018.689017. DOI: 10.1101/2025.11.18.689017.

(53) Lundby, A.; Secher, A.; Lage, K.; Nordsborg, N. B.; Dmytriyev, A.; Lundby, C.; Olsen, J. V. Quantitative maps of protein phosphorylation sites across 14 different rat organs and tissues. Nat. Commun 2012, 3, 876. DOI: 10.1038/ncomms1871.

(54) Huttlin, E. L.; Jedrychowski, M. P.; Elias, J. E.; Goswami, T.; Rad, R.; Beausoleil, S. A.; Villen, J.; Haas, W.; Sowa, M. E.; Gygi, S. P. A tissue-specific atlas of mouse protein phosphorylation and expression. Cell 2010, 143 (7), 1174–1189. DOI: 10.1016/j.cell.2010.12.001.

(55) Bai, Y.; Tolles, J.; Cheng, H.; Siddiqui, S.; Gopinath, A.; Pectasides, E.; Camp, R. L.; Rimm, D. L.; Molinaro, A. M. Quantitative assessment shows loss of antigenic epitopes as a function of pre-analytic variables. Lab. Invest 2011, 91 (8), 1253–1261. DOI: DOI: 10.1038/labinvest.2011.75.

(56) Pinhel, I. F.; Macneill, F. A.; Hills, M. J.; Salter, J.; Detre, S.; A’Hern, R.; Nerurkar, A.; Osin, P.; Smith, I. E.; Dowsett, M. Extreme loss of immunoreactive p-Akt and p-Erk1/2 during routine fixation of primary breast cancer. Breast. Cancer. Res 2010, 12 (5), R76. DOI: DOI: 10.1186/bcr2719.

(57) Yaron-Barir, T. M.; Joughin, B. A.; Huntsman, E. M.; Kerelsky, A.; Cizin, D. M.; Cohen, B. M.; Regev, A.; Song, J.; Vasan, N.; Lin, T.-Y.;, et al. The intrinsic substrate specificity of the human tyrosine kinome. Nature 2024, 629 (8014), 1174–1181. DOI: 10.1038/s41586-024-07407-y.

(58) Johnson, J. L.; Yaron, T. M.; Huntsman, E. M.; Kerelsky, A.; Song, J.; Regev, A.; Lin, T.-Y.; Liberatore, K.; Cizin, D. M.; Cohen, B. M.;, et al. An atlas of substrate specificities for the human serine/threonine kinome. Nature 2023, 613 (7945), 759–766. DOI: 10.1038/s41586-022-05575-3.

(59) Keshishian, H.; McDonald, E. R.; Mundt, F.; Melanson, R.; Krug, K.; Porter, D. A.; Wallace, L.; Forestier, D.; Rabasha, B.; Marlow, S. E.;, et al. A highly multiplexed quantitative phosphosite assay for biology and preclinical studies. Mol Syst Biol 2021, 17 (9), MSB202010156. DOI: 10.15252/msb.202010156.

(60) Wang, D.; Eraslan, B.; Wieland, T.; Hallström, B.; Hopf, T.; Zolg, D. P.; Zecha, J.; Asplund, A.; Li, L. h.; Meng, C.;, et al. A deep proteome and transcriptome abundance atlas of 29 healthy human tissues. Mol Syst Biol 2019, 15 (2), MSB188503. DOI: 10.15252/msb.20188503.

(61) Uhlén, M.; Fagerberg, L.; Hallström, B. M.; Lindskog, C.; Oksvold, P.; Mardinoglu, A.; Sivertsson, Å.; Kampf, C.; Sjöstedt, E.; Asplund, A.;, et al. Tissue-based map of the human proteome. Science 2015, 347 (6220), 1260419. DOI: 10.1126/science.1260419.

(62) Thul, P. J.; Åkesson, L.; Wiking, M.; Mahdessian, D.; Geladaki, A.; Ait Blal, H.; Alm, T.; Asplund, A.; Björk, L.; Breckels, L. M.;, et al. A subcellular map of the human proteome. Science 2017, 356 (6340), eaal3321. DOI: 10.1126/science.aal3321 (accessed 2026/07/02).

(63) Karlsson, M.; Zhang, C.; Méar, L.; Zhong, W.; Digre, A.; Katona, B.; Sjöstedt, E.; Butler, L.; Odeberg, J.; Dusart, P.;, et al. A single–cell type transcriptomics map of human tissues. Science Advances 2021, 7 (31), eabh2169. DOI: 10.1126/sciadv.abh2169 (accessed 2026/07/02).

(64) Uhlen, M.; Zhang, C.; Lee, S.; Sjöstedt, E.; Fagerberg, L.; Bidkhori, G.; Benfeitas, R.; Arif, M.; Liu, Z.; Edfors, F.;, et al. A pathology atlas of the human cancer transcriptome. Science 2017, 357 (6352), eaan2507. DOI: 10.1126/science.aan2507 (accessed 2026/07/02).

